# Methanogenesis inhibition remodels microbial fermentation and stimulates acetogenesis in ruminants

**DOI:** 10.1101/2024.08.15.608071

**Authors:** Gaofeng Ni, Nicola Walker, André Fischer, René T. Stemmler, Oliver Schmidt, Surbhi Jain, Marion Jespersen, Rhys Grinter, Min Wang, Phillip B. Pope, Volker Müller, Mick Watson, Emiel Ver Loren van Themaat, Maik Kindermann, Chris Greening

**Affiliations:** Department of Microbiology, Biomedicine Discovery Institute, Monash University, Melbourne, Australia; DSM Nutritional Products, Animal Nutrition and Health, Kaiseraugst, Switzerland; dsm-firmenich, Beauty & Care, Kaiseraugst, Switzerland; dsm-firmenich, Chemical & Process Sciences, Kaiseraugst, Switzerland; UIT The Arctic University of Norway, Tromsø, Norway; Key Laboratory for Agro-Ecological Processes in Subtropical Region, Institute of Subtropical Agriculture, Chinese Academy of Sciences, Hunan, China; University of Chinese Academy of Sciences, Beijing, China; Centre for Microbiome Research, Queensland University of Technology, Brisbane, Australia; Faculty of Chemistry, Biotechnology and Food Science, Norwegian University of Life Sciences, Ås, Norway; Department of Molecular Microbiology and Bioenergetics, Institute of Molecular Biosciences, Johann Wolfgang Goethe University, Frankfurt, Germany; Data Science, Science & Research, dsm-firmenich, Delft, The Netherlands

## Abstract

Rumen microbiota enable ruminants to grow on fibrous plant materials but also produce methane, driving 5% of global greenhouse gas emissions and leading to a loss of gross energy content. Methanogenesis inhibitors such as 3-nitrooxypropanol (3-NOP) decrease methane emissions in ruminants when supplemented in feed. Yet we lack a system-wide, species-resolved understanding of how the rumen microbiota remodels following inhibition and how this influences animal production. Here, we conducted a large-scale trial with 51 dairy calves to analyse microbiota responses to 3-NOP, pairing host performance, emissions, and nutritional profiles with genome-resolved metagenomic and metatranscriptomic data. 3-NOP supplementation decreased methane emissions by an average of 62%, modulated short-chain fatty acid and H_2_ levels, and did not affect dietary intake or animal performance. We created a rumen microbial genome catalogue with an unprecedented mapping rate. We observed a strong reduction of methanogens and stimulation of reductive acetogens, primarily novel uncultivated lineages such as *Candidatus* Faecousia. However, there was a shift in major fermentative communities away from acetate production in response to hydrogen gas accumulation. Thus, the divergent responses of the fermentative and hydrogenotrophic communities limit potential productivity gains from methane reduction. Reporting one of the largest reductions in methane emissions in a field trial to date, this study links ruminant greenhouse gas emissions and productivity to specific microbial species. These findings also emphasise the importance of microbiota-wide analysis for optimising methane mitigation strategies and identify promising strategies to simultaneously reduce emissions while increasing animal production.

**Significance Statement:** One strategy to increase the sustainability and productivity of livestock production is to modulate ruminant microbiota to produce absorbable nutrients rather than the potent greenhouse gas methane. Previous studies show supplementing feed with methanogenesis inhibitors such as 3-nitrooxypropanol reduces methane emissions, but also leads to inconsistent productivity gains. Here we report a definitive field trial, combining animal data, meta-omics, and structural modelling, to resolve the key microbes and pathways controlling nutrient and methane production in ruminants. We show that shifts in composition and gene expression of hydrogen-cycling microbes reduce emissions but limit productivity gains. These findings offer insights at unprecedented resolution, while the data and analytical framework provide valuable resources to develop solutions to enhance livestock productivity and sustainability.

## Introduction

Methane (CH_4_) mitigation offers an opportunity to enhance the sustainability and productivity of the livestock sector. Globally, ruminant enteric emissions were responsible for 27 to 31 % of anthropogenic CH_4_ emissions in 2020^1,2^. Moreover, these CH_4_ emissions represent an average of 6% loss of gross energy intake from feed^3^. Ruminants can digest fibrous feed because of their unique microbiota in their rumen. Anaerobic bacteria, fungi, and protists hydrolyse and ferment carbohydrates into short-chain fatty acids (SCFAs), carbon dioxide (CO_2_), and hydrogen (H_2_)^4^. SCFAs, such as acetate, butyrate, and propionate^5^, are primary energy sources for the host. In contrast, H_2_ is primarily consumed by hydrogenotrophic microorganisms, enabling digestion and fermentation to remain thermodynamically favorable^6,7^. The dominant hydrogenotrophs in the rumen are methanogenic archaea, including those that use H_2_ as an energy source and electron donor to reduce CO_2_, formate, or methanol to CH_4_, and are ubiquitous in the gastrointestinal tracts of diverse ruminants. While other hydrogenotrophs also inhabit the rumen, including acetogens and sulfate-, nitrate-,, and fumarate-reducing microorganisms, they are typically less abundant and active than methanogens^6^. Redirecting ruminal H_2_ flow away from methanogenesis towards alternative hydrogenotrophic pathways such as acetogenesis is one of the most promising strategies to mitigate enteric CH_4_ emissions while maintaining ruminant health and productivity^6,8^. However, the H_2_ threshold for reductive acetogenesis (430 - 950 ppm) is typically much higher than for methanogenesis (25 - 100 ppm), putting acetogens at a disadvantage when competing with methanogens for H_2_ in the rumen^9^. Currently, we lack a system-wide and species-resolved understanding of what controls ruminant electron flow, including the processes and microbes mediating H_2_ production and consumption. Building an evidence base on these aspects is critical to inform and assess intervention strategies to redirect electron flows away from methane and into production.

Among the most promising strategies to reduce CH_4_ emissions are to apply direct-acting methanogenesis inhibitors^6,10–13^. One such inhibitor, 3-nitrooxypropanol (3-NOP; marketed as Bovaer^®^, dsm-firmenich, Switzerland), has been shown to be a potent and specific inhibitor of methanogenesis in the rumen. Field trials show 3-NOP effectively reduces CH_4_ emissions in beef cattle (27 to 90% reduction)^14–16^ and dairy cows (23 to 60% reduction)^17–20^, though there are considerable variations in efficacy between trials, driven mainly by diet composition, inhibitor dosage, and type of animal (e.g., dairy or beef cattle). The mode-of-action of 3-NOP has been elucidated in the model organism *Methanothermobacter marburgensis*^12^: 3-NOP inactivates the key enzyme for methanogenesis, methyl-CoM reductase (MCR), by binding to its active site and oxidising the nickel ion (Ni^+^) within the cofactor F_430_, which is active only in the +1 oxidation state^12^. However, its efficacy varies between methanogen species, with both culture-based^12^ and culture-independent^17^ studies showing more potent inhibition of hydrogenotrophic methanogens (e.g., *Methanobrevibacter*, *Methanobacterium*) than methylotrophic or metabolically flexible methanogens (e.g., *Methanosarcina*, *Methanomethylophilus*). While CH_4_ production results in loss of gross energy for ruminants, supplementation of methanogenesis inhibitors such as 3-NOP leads to variable and sometimes negligible productivity gains across field trials^11,20–23^ for reasons that have yet to be resolved.

In this context, it is unclear how the wider rumen ecosystem responds to supplementation of methanogenesis inhibitors and whether this explains limited production gains. When rumen methanogenesis is inhibited by 3-NOP, *in vivo* H_2_ emissions increase by approximately four-fold^11,23,24^. Elevated rumen H_2_ partial pressures may lead to cascading effects in rumen metabolism, given that most rumen microorganisms encode hydrogenases^6^. Given acetogens are typically outcompeted by methanogens due to their higher H_2_ threshold^6,9^, the accumulation of rumen H_2_ caused by the inhibition of methanogenesis^11,23,24^ may provide an opportunity for hydrogenotrophic acetogens to flourish, though has not been reported to date. In contrast, high *in situ* H_2_ concentrations limit the possibility for fermenters to dispose of excess reductants as H_2_, and as a consequence they may need to shift the fermentation profile towards the production of more reduced organic compounds such as ethanol instead of acetate, as seen through culture-based studies of *Ruminococcus albus*^6,25^. The extent to which these predicted processes occur in the rumen and impact production gains is unknown.

Here we address these knowledge gaps by performing a definitive study of microbiome-wide responses to supplementation of methanogenesis inhibitors. We investigated whether administration of 3-NOP during calf development stimulates the development of a low methane-emitting community^26,27^. To do so, we conducted a large field trial of 3-NOP supplementation by combining animal performance, nutrition, and emissions data with species-resolved profiling of the microbes and pathways responsible for electron flows and H_2_ cycling. These results show that, while 3-NOP potently inhibits methanogenesis and stimulates acetogens, corresponding community and metabolic shifts in fermenter communities limit productivity gains. Collectively, these findings suggest that interventions to reduce emissions and increase productivity are best achieved by considering the response of the entire microbiota to the direct and indirect effects of methanogenesis inhibition.

## Results and Discussion

### 3-nitrooxypropanol inhibits methane production, which leads to the modulation of SCFA and H_2_ levels

In a field trial involving 51 Holstein X Jersey calves for 99 days, 17 animals were assigned to three 3-NOP dosage groups: control (no 3-NOP), low (target dose rate of 150 mg 3-NOP / kg dry matter intake, DMI, per day), and high (target dose rate of 250 mg 3-NOP / kg DMI per day). Mean CH_4_ emissions were reduced significantly by 60.0% (*p* = 1.7 × 10^-5^) and 62.3% (*p* = 1.5 × 10^-5^) in low and high 3-NOP dosage groups, respectively, compared to the no intake control (**Fig. 1a**). The observed reduction in CH_4_ emissions is one of the largest reductions for *in vivo* trials to date. Meanwhile, no significant changes were observed in body weight gain, milk and meal intake, and feed energy efficiency (*p* > 0.05) (**Fig. 1b**), which is consistent with previous studies with adult beef and dairy cattles^11,21–23,28,29^. This indicates that 3-NOP supplementation significantly reduces CH_4_ emissions, but does not affect dietary intake and growth performance of calves.

**Figure 1:**
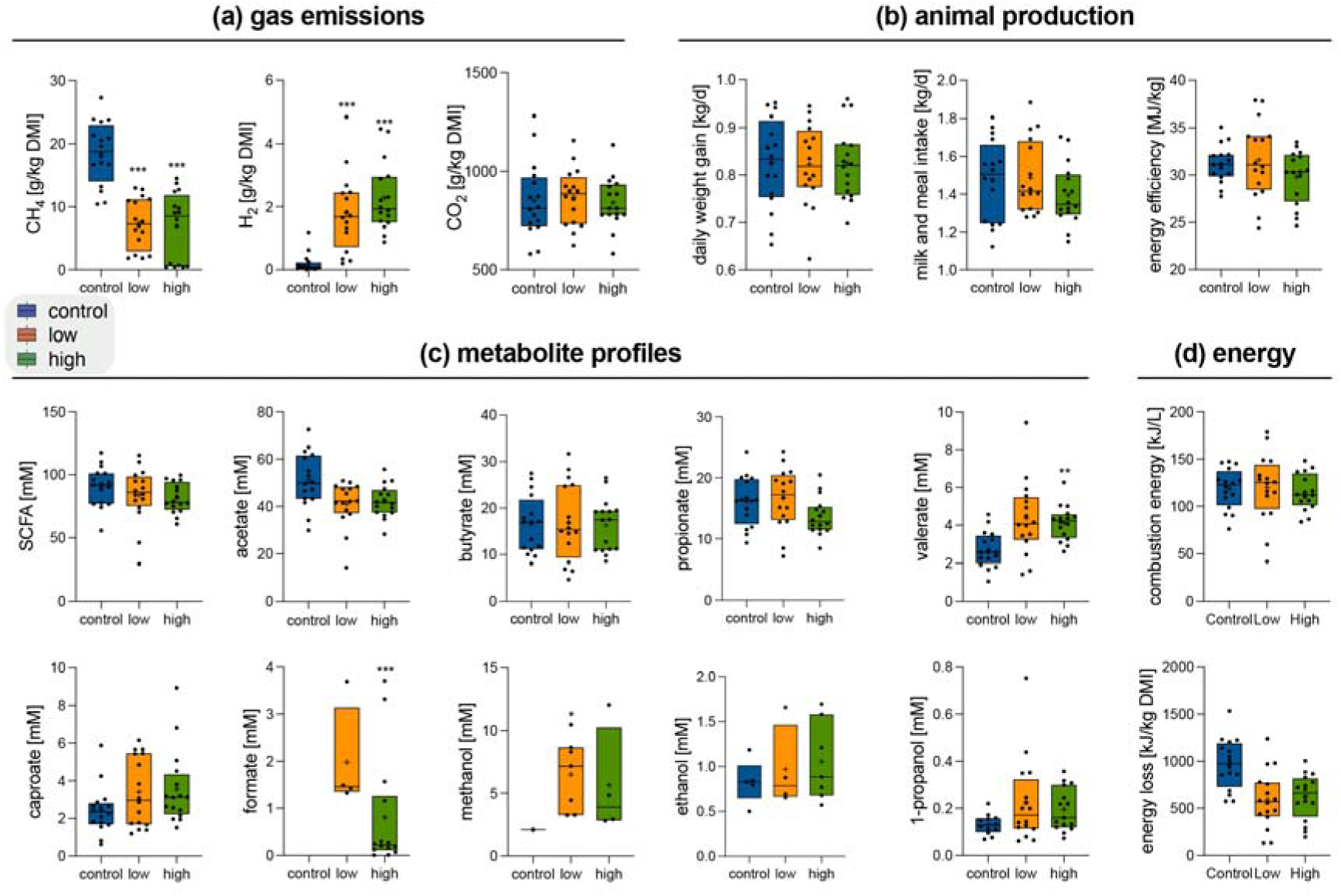
Effects of 3-NOP on gas emissions, animal production, ruminal metabolites, and energy states. CH_4_, H_2_, and CO_2_ emissions calculated as gram per kilogram of dry matter intake (**a**), animal diet and performance metrics (**b**), concentrations of ruminal short chain fatty acids (SCFAs) and alcohols (**c**), and combustion energy of metabolites and energy loss from CH_4_ and H_2_ emissions (**d**). The statistical significance of differences between groups (low versus control and high versus control) for each parameter was assessed using the Kruskal–Wallis method with a false discovery rate correction (*p < 0.05, **p < 0.01, ***p < 0.001). Treatment groups are differentiated by colour (red, green and blue for no, low and high 3-NOP, respectively) in all box plots.

Following 3-NOP supplementation, H_2_ emissions increased by 9.0-fold (low dose, *p* = 2.6 × 10^-5^) and 11.5-fold (high dose, *p* = 6.1 × 10^-6^) (**Fig. 1a**). This suggests that other hydrogenotrophs in the rumen do not fully consume the H_2_ that would otherwise be metabolised by methanogens. Another methanogenesis substrate, formate, was only detectable in the low and high groups. This likely reflects the suppression of methanogens that express formate dehydrogenases (*fdhA*) to use formate as a substrate^30,31^, though may also reflect shifts in the activities of fermenters. Methanol levels also increased 2.9- and 5.2- fold in the low and high groups, respectively, likely resulting from the inhibition of methylotrophic methanogenesis^32^. While the overall concentrations of SCFAs did not significantly change with 3-NOP supplementation, there were modest but significant changes in both the absolute concentrations and molar proportions of specific fatty acids; notably, acetate concentrations decreased (1.24-fold low 3-NOP, 1.21-fold high 3-NOP), whereas both caproate (1.41-, 1.47-fold) and valerate (1.61-, 1.52-fold) increased, with no significant effects on propionate or butyrate levels (**Fig. 1c**). The combustion energy of rumen SCFAs, reflecting the total amount of chemical energy available for the ruminant, did not significantly change across the three experimental groups (**Fig. 1d**). It should be cautioned that the concentrations reported are for circulating metabolites in the rumen, but SCFAs and alcohols are absorbed by rumen epithelium at rapid and likely variable rates, in addition to being consumed by other rumen microbes^5,33–38^.

To determine why productivity gains were not observed, we compared the energy and reductant loss between the three groups (**Fig. S1**). Calves of the no, low, and high 3-NOP treatment groups lost on average 970, 591, and 622 kJ / kg DMI per day due to the emissions of H_2_ and CH_4_, respectively (**Fig. 1d)**. Thus, the energy loss due to H_2_ and CH_4_ emissions was reduced by 39.1% and 35.9% in low and high 3-NOP groups, respectively (**Fig. S1b**), relative to untreated calves. Considering that approximately 6% of gross energy intake is lost due to CH_4_ emissions^3^ and loss due to H_2_ emissions is negligible in untreated cows (**Fig. S1b**), 3-NOP administration in this trial is predicted to increase rumen performance parameters such as body weight gain or energy efficiency by approximately 2%. However, no significant differences in animal parameters were observed in our study, likely reflecting the sample size is insufficient to capture these modest potential gains, especially given the high variability in performance metrics between individual cows.

### 3-nitrooxypropanol addition remodels metabolic profiles of rumen microbiota

Paired metagenomic and metatranscriptomic short reads of the rumen microbial communities were analysed to determine the structure and function of the rumen microbiota. 3-NOP supplementation did not significantly influence the richness and composition of the microbial communities, as there was a lack of clustering between groups in beta diversity analyses (*p* = 0.479) (**Fig. 2a, b**). While 3-NOP did not affect the abundance of the dominant bacterial phyla, as expected, it broadly reduced the abundance of the methanogenic archaeal phyla Methanobacteriota and Thermoplasmatota. Most notably, Methanobacteriota decreased in relative abundance by 2.7- (low dose, *p* = 2.5 × 10^-5^) and 4.7-fold (high dose, *p* = 1.5 × 10^-5^) based on mapping against metagenomic reads. Concordantly, the gene expression of Methanobacteriota decreased by 2.0-fold at both doses based on the mapping of metatranscriptomic reads (**Fig. 2c**). This finding agrees with previous studies showing that Methanobacteriota is more affected by 3-NOP than other methanogen lineages^17,39^. The abundance changes of these groups corroborate with the changes of the functional profiles of the microbiota. For instance, a significant and dose-dependent reduction was observed in the gene encoding the catalytic subunits of the methane-producing enzyme methyl-CoM reductase (*mcrA*; by 3.5-fold <*p* = 2.0 × 10^-5^> and 6.5-fold <*p* = 6.9 × 10^-6^> under low and high 3-NOP), as well as the catalytic subunits of methanogen-type hydrogenases (groups 3a, 3c, 4h, 4i [NiFe]-hydrogenases, and [Fe]-hydrogenase) (**Fig. 2d, e**). Similar trends were observed based on the metatranscriptomic data, with the expression of the methyl-CoM reductase declining below that of the enzymes mediating acetogenesis, sulfate reduction, and nitrate reduction following 3-NOP supplementation (**Fig. 2f**). These findings support previous observations that 3-NOP potently and specifically inhibits the growth and activity of methanogens^17,40^.

**Figure 2:**
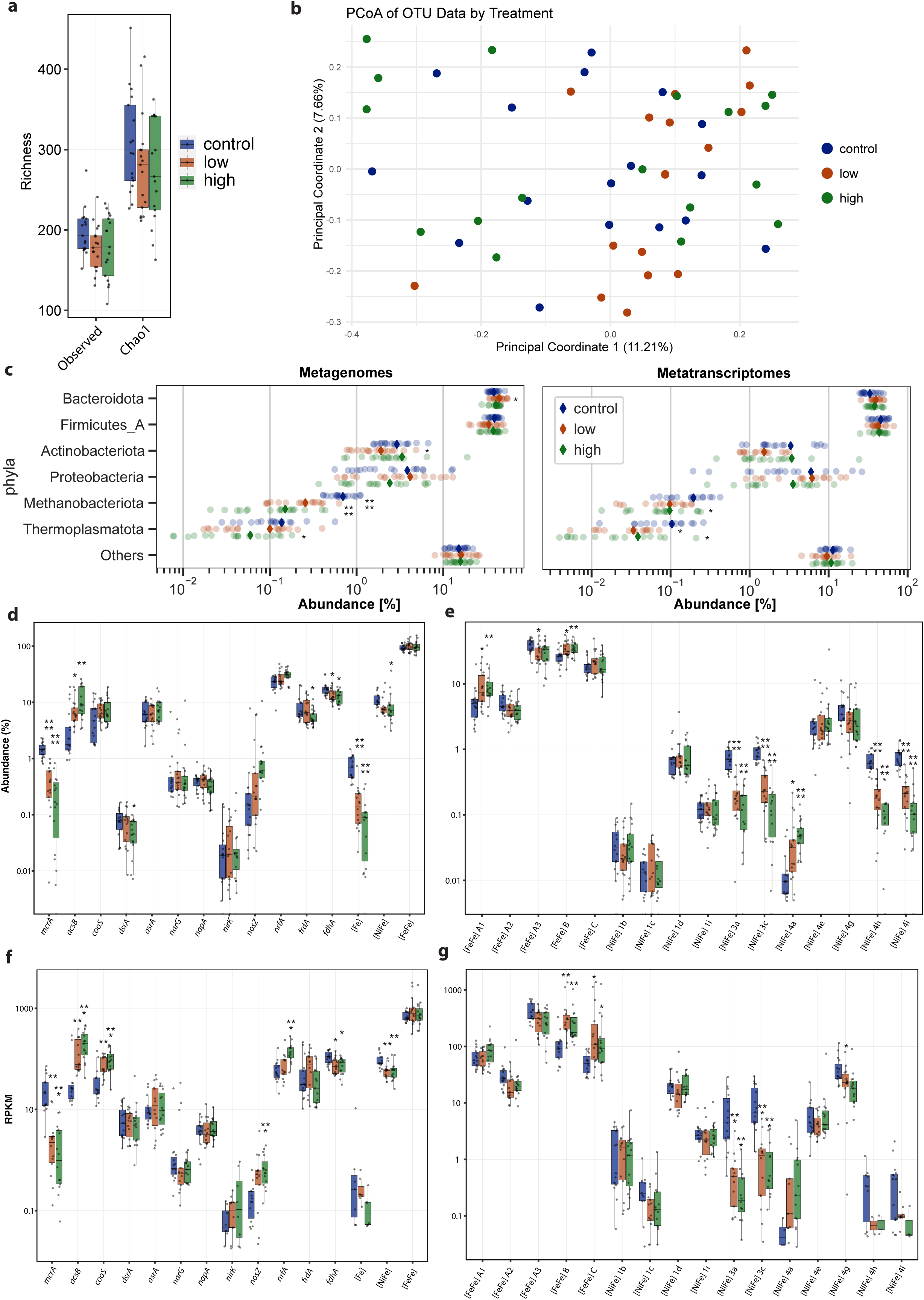
Effect of 3-NOP on composition and function of rumen microbiota. (**a**) Boxplots displaying alpha diversity, measured as observed and estimated richness (Chao1), across three groups. Significance testing was conducted using the Kruskal-Wallis test with a false discovery rate correction; no statistically significant differences were found. (**b**) Principal coordinates analysis (PCoA) plot illustrating beta diversity and coloured by group, using an abundance-based distance matrix calculated with the Bray-Curtis method. Significant differences in community structure between groups were tested using one-way PERMANOVA with 999 permutations. (**c**) Scatter plots showing the distribution of abundance and activity of taxa aggregated at the phylum level. The abundance of each taxonomic group was inferred from metagenomic (left) and metatranscriptomic short reads (right). Diamonds indicate mean abundance values. Statistical significance of differences between groups (low versus control and high versus control) for relative abundance was assessed using Mann–Whitney *U* test (\**p* < 0.05, \*\**p* < 0.01, \*\*\**p* < 0.001, \*\*\*\**p* < 0.0001). The abundance (**d**) and expression (**f**) of marker genes for reductive processes; the abundance (**e**) and expression of hydrogenase sub-groups (**g**). The statistical significance of differences between groups (low versus control and high versus control) for each gene was tested using the Kruskal–Wallis test with a false discovery rate correction using the Benjamini-Hochberg procedure (\**p* < 0.05, \*\**p* < 0.01, \*\*\**p* < 0.001, and \*\*\*\**p* < 0.0001). Treatment groups are differentiated by colour (red, green and blue for no, low and high 3-NOP, respectively) in all box plots.

We expanded our analysis to evaluate the abundance and expression of other marker genes mediating other H_2_ cycling processes to better understand the modulation of alternative H_2_ sinks in the rumen upon 3-NOP inhibition. Communities involved in H_2_ production shifted, as evidenced by the increased abundance and expression of ferredoxin-dependent, monomeric group A1 and B [FeFe]-hydrogenases that are energetically advantageous under high H_2_ concentrations, at the expense of trimeric electron bifurcating or confurcating group A3 [FeFe]-hydrogenases (**Fig. 2e, g**)^6^. The marker genes and transcripts for alternative respiratory processes were also detected, including for fumarate (*frdA*), nitrate (*narG*, *napA*), and sulfite (*dsrA*, *asrA*) reduction^41,42^. 3-NOP supplementation appeared to have minor effects on these pathways. In contrast, a profound increase in the abundance and expression of acetogens was observed, based on the genes encoding the acetyl-CoA synthase / carbon monoxide dehydrogenase complex central to Wood-Ljungdahl pathway (WLP) (*acsB* by 1.4- and 2.7-fold, *cooS* by 2.7- and 4.0-fold at low and high 3-NOP doses respectively). Transcripts for acetogenesis were higher than those of any other known ruminal hydrogen sinks (42-, 28-, 1.5-, 297-, and 5.6-fold higher than *mcrA*, *dsrA*, *nrfA*, *nosZ*, and *frdA*, respectively) (**Fig**. **2f****; Dataset S1**). Together, these data suggest that reductive acetogenesis becomes the dominant H_2_ consumption pathway following 3-NOP supplementation.

### 3-NOP strongly inhibits methanogens from multiple genera by inactivating the methyl-CoM reductase

We magnified our focus to the species level to better understand how the metabolism of individual microbes influences ruminant emissions and productivity. To do so, we mapped metagenomic and metatranscriptomic reads from the field trial to a comprehensive collection of rumen genomes. This compendium was compiled by constructing metagenome-assembled genomes (MAGs) from three field trials of 3-NOP, including this trial, and consolidating them with MAGs and isolate genomes reported from eight other datasets^43–50^. From the dereplication of the 27,884 resultant MAGs and isolate genomes, 11,909 medium-to high-quality, species-resolved genomes were obtained, which includes 3,003 rumen microbial species that were present in field trial animals (**Dataset S2**). This genome collection provides unprecedented species-level representations of ruminant microbiota, mapping to 76.7% and 80.5% of the metagenomic and metatranscriptomic reads from the field trial, respectively.

These analyses revealed the trial animals harbour 49 methanogen species, spanning four genera: *Methanobrevibacter*, *Methanobrevibacter*_A, and *Methanosphaera* from the Methanobacteriota phylum and *Methanomethylophilus* from the Thermoplasmatota phylum (**Fig. 3a**). 3-NOP broadly affected these methanogens, significantly reducing their abundance by an average of 3.7- and 6.3-fold at low and high 3-NOP doses, respectively (**Fig. 3b**). The most pronounced inhibition was observed in *Methanobrevibacter*, with a 15-fold decrease in its overall abundance and a 748-fold decrease in the most abundant *Methanobrevibacter* MAG at high doses. The weakest inhibition was observed in *Methanomethylophilus*, with decreases of 1.9-fold and 2.8-fold at low and high doses, respectively. Genome-resolved metatranscriptomic analysis revealed variable and generally low gene expression among the methanogen species across all groups (**Fig. 3b**). In line with metagenomic findings, 3-NOP inhibited methanogen gene expression, showing the strongest in *Methanobrevibacter* (up to 100% reduction) and to a less degree in *Methanomethylophilus* (76% and 73% reduction at low and high 3-NOP doses, respectively). Differential expression analysis shows potent decrease in expression of the methanogenesis pathway in *Methanobrevibacter* (**Fig. S2**). Residual CH_4_ emissions may originate from the activities of the methylotrophic *Methanomethylophilus* populations, and potentially due to the occurrence of methanogens within other components of the gastrointestinal tract such as the jejunum, ileum, and cecum^43,51^. The increased H_2_ and formate accumulation (**Fig. 1a, b**) likely reflects the inhibition of methanogens encoding hydrogenases and formate dehydrogenases (lineage-specific variations shown in **Fig. 3b**), in line with the significant decrease in abundance and expression of methanogen-type hydrogenases and *fdhA* at the community level (**Fig. 2d, f**).

**Figure 3:**
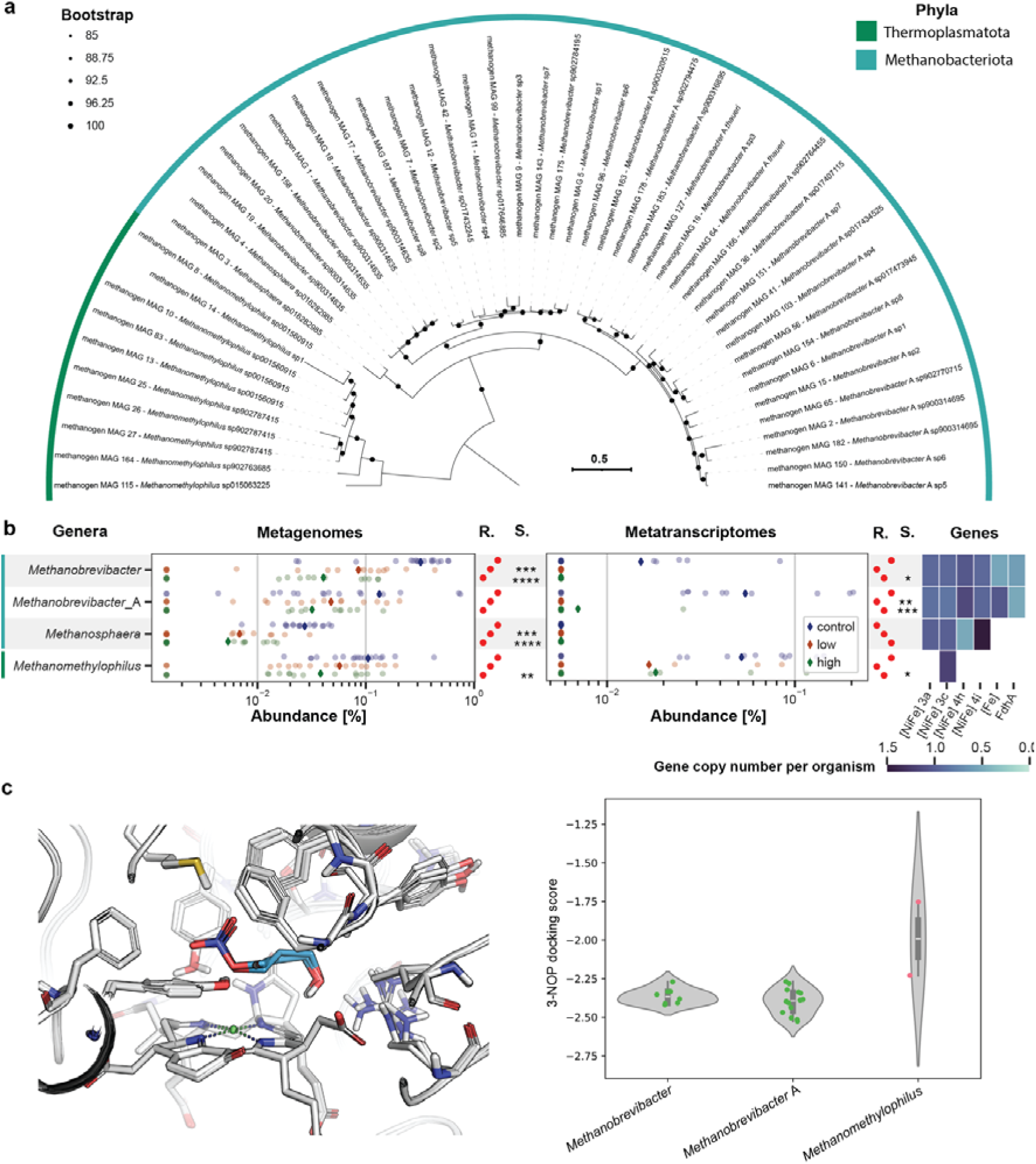
Effects on 3-NOP on methanogen populations in the rumen. (**a**) Phylogenomic tree of the 49 methanogen MAGs used in this study, inferred through the maximum likelihood method (VT+F+G4) and 1,000 bootstrap iterations. The tree was rooted at the midpoint. (**b**) Scatter plots showing the abundance (left) and activity (right) of four methanogen genera, analysed through mapping against metagenomes and metatranscriptomes. Diamonds indicate mean abundance values, and a red dot plot shows rank abundance (indicated as “R.”) from low (left) to high (right). Statistical significance (indicated as “S.”) of differences between groups (low versus control and high versus control) for MAGs was assessed using Mann–Whitney *U* test (*p < 0.05, **p < 0.01, ***p < 0.001, ****p < 0.0001). The heatmap shows the average copy number of the group 3a, 3c, 4h, 4i [NiFe]-hydrogenases and [Fe]-hydrogenase encoded by these organisms. (**c**) Left: a close-up view of 3-NOP binding based on docking calculations in the active site of superimposed predicted structures of MCR. MCRs from the most abundant species are shown in cartoon, with one each from *Methanobrevibacter*_A, *Methanobrevibacter*, *Methanosphaera*, and *Methanomethylophilus*. The active site residues are shown in stick representation with carbon, nitrogen, oxygen, and sulfur atoms coloured in white (cyan for 3-NOP), blue, red, and yellow, respectively. The nickel ion of the F_430_ cofactor is shown as a green sphere. Right: A violin plot depicting docking scores that were computed using the Glide SP docking protocol.

To rationalise these observations, structural models of methyl-CoM reductase (MCR) from the 49 species identified in this study were generated and refined using AlphaFold2, and then 3-NOP was docked(**Fig. 3c, S3, Dataset S3**). Based on the docking predictions, 3-NOP fits well into the active site of all MCR structures, explaining the broad inhibition of methanogenesis across the methanogen lineages detected in the dataset. The active sites and binding tunnels of the enzyme are highly conserved, with just three amino acid substitutions across the dataset, at positions that would minimally affect 3-NOP binding. Thus, differences in MCR structures cannot sufficiently explain the variable efficacy of 3-NOP on different methanogens. Instead, lineage specific activation systems may contribute to the variation in 3-NOP inhibition efficacies^52,53^. Overall, these data confirm that 3-NOP is a potent methanogenesis inhibitor, though it varies in efficacy between methanogen species and genera for yet unresolved reasons.

### Fermentation shifts away from acetate production in response to H_2_ accumulation

To resolve whether 3-NOP supplements influence fermentation pathways, we performed a pathway-centric analysis based on the fold change of transcripts from fermentative species (**Fig. 4a**). There was an overall decrease in transcripts for fermentative acetate formation, yet an increase in WLP transcripts relative to control samples, signifying the upregulation of acetogenesis. Given the measured decrease in total acetate concentrations in rumen samples (**Fig. 1c**), our results suggest that decreased acetate fermentation has a stronger effect than increased hydrogenotrophic acetogenesis, thereby negating potential productivity gains from methanogenesis inhibition. We also observed upregulation of butyrate and ethanol production pathways, particularly at high 3-NOP dosage. Propionate production pathways overall were not affected by 3-NOP, with minor downregulation of the acrylate pathway and minor upregulation of the succinate pathway. 3-NOP supplementation stimulated reverse beta oxidation, leading to the production of valerate and caproate^54–56^, as confirmed by increased concentrations of these SCFAs (**Fig. 1c**). However, the total metabolite profile did not significantly change, in line with the unchanged growth parameters in calves (**Fig. 1b, d**). These observations were further confirmed through analysis of fermentation marker genes by short read analysis (**Fig. S4a**).

**Figure 4:**
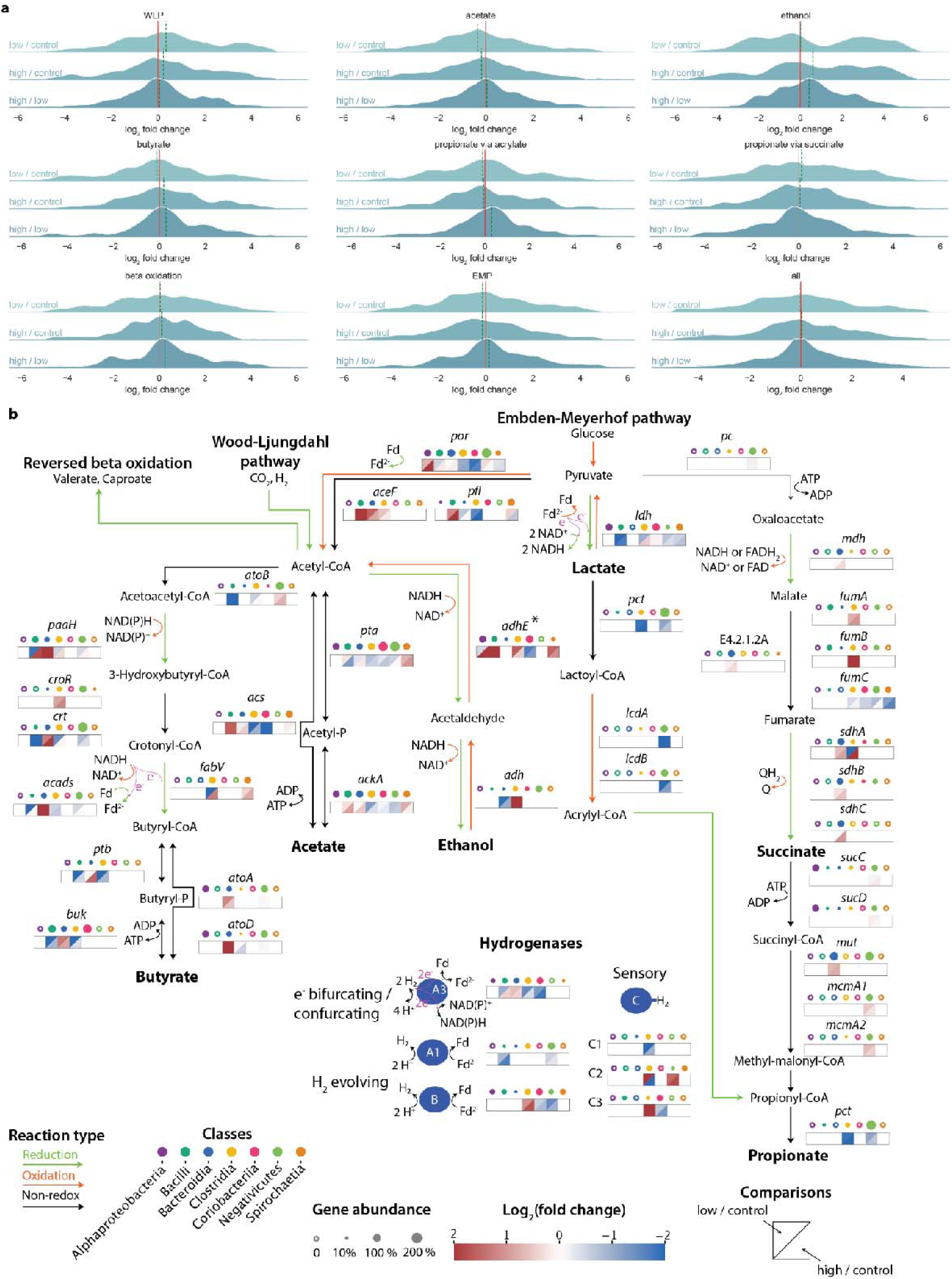
Effects on 3-NOP on the processes and mediators of ruminal fermentation. (**a**) Comparative metatranscriptomic analysis of rumen SCFA production pathways based on the mapping of metatranscriptomic reads onto each rumen MAG. The comparisons are separated into high versus control, low versus control, and high versus low, differentiated by light to darker shades of green. The SCFA metabolism categories include Wood-Ljungdahl Pathway (WLP), conversion from pyruvate to acetate, conversion of acetyl-CoA to ethanol, conversion of acetyl-CoA to butyrate, conversion from pyruvate to propionate via acrylate, conversion from pyruvate to propionate via succinate, beta oxidation (KEGG module M00087), conversion of glucose to pyruvate (Embden-Meyerhof pathway, EMP), and finally all identified transcripts. The plot area represents the normalised fold change (log_2_) of transcripts per category, calculated using the kernel density estimate (KDE) method implemented in the Python package seaborn. The red lines indicate no fold change (log_2_ = 0), while the median value for each pathway is indicated by green dashed lines. (**b**) Fermentation pathways of the 30 most abundant genera predicted to mediate fermentation. The abundance of each gene was calculated as copy number percent aggregated by the class-level taxonomy of the microorganisms encoding these genes, differentiated by colour. The heatmap indicates the expression fold change (low / control at the top left corner and high / control at the bottom right corner) for specific genes that was averaged at the class level. *The bifunctional acetaldehyde-alcohol dehydrogenase (*adhE*) converts acetyl-CoA to ethanol via the intermediate acetaldehyde^59^. The full list of gene abbreviations is available in **Dataset S5.**

To determine the microbes responsible for shifts in fermentation pathways, we identified the 30 most abundant fermentative genera that encode group A1, A3, and B [FeFe]-hydrogenases thought to be primarily responsible for H_2_ production in the rumen^6^, and investigated whether they encoded and differentially expressed pathways for acetate, butyrate, and propionate production (**Fig. 4b**). Notably, 11 of these genera currently lack cultivated representatives. In accordance with literature^4^, diverse lineages are capable of fermentative acetate production. Ethanol production capabilities are also widespread, and the associated genes were upregulated in Clostridia (*Massiliimalia*, *Ruminococcus*_E, UBA2862, and SFMI01), Bacilli (*Sharpea*), and Bacteroidia (*Sodaliphilus*) in response to 3-NOP supplementation. Butyrate production capabilities were conserved within not only Clostridia, in line with previous reports^4^, but also in species affiliating with Bacilli, Bacteroidia, and Negativicutes. The stimulation of butyrate and propionate production in response to 3-NOP supplementation was predominantly observed in Bacteroidia, the latter primarily via the succinate pathway. Among the 30 most abundant fermenters, Negativicutes was the only class to encode the full acrylate pathway, the expression of which was downregulated in response to 3-NOP supplementation.

Upon 3-NOP supplementation, various clostridial genera downregulate the group A3 [FeFe]-hydrogenases in favour of the group A1 or B [FeFe]-hydrogenases. The trimeric A3 enzymes confurcate electrons from NADH and ferredoxin to H_2_, whereas the group A1 and B [FeFe]-hydrogenases as monomeric ferredoxin-dependent H_2_-producing enzymes are less inhibited by H_2_ accumulation^57,58^. These genera also express putative sensory C3 [FeFe] hydrogenases that are typically co-transcribed with H_2_-evolving hydrogenases^58^ (**Fig. 4b**). Multiple genera shift their hydrogenase expression as such following 3-NOP supplementation, including *Ruminococcus*_E, *Sodaliphilus*, W0P33-017, UBA3282, and *Massiliimalia*. This shift in expression is likely a direct response to H_2_ accumulation, as probably sensed by group C [FeFe] hydrogenases, resulting in upregulation of the monomeric hydrogenases and alcohol dehydrogenase at the expense of the electron-confurcating hydrogenase. This enables fermentation to continue at high H_2_ partial pressures, resulting in less ATP and lower H_2_ production, in line with our culture-based studies with *Ruminococcus albus* 7^6,25^. Given the high level of abundance and expression of sensory hydrogenases (**Fig. 2e, g**), this finding suggests H_2_ cycling is tightly intertwined with fermentation and that hydrogen sensing is a general mechanism regulating hydrogenase expression in ruminants^4^. Regulation of hydrogenase activity through hydrogen sensing may therefore be a crucial aspect of microbial responses to 3-NOP supplementation.

In addition to differential regulation of hydrogenases, there was also evidence of differential regulation of SCFA production pathways (**Dataset S4**). In general, fermentative microorganisms that were stimulated appear to regulate multiple fermentation routes that can help them adapt to thermodynamic challenges caused by H_2_ accumulation. For instance, *Sodaliphilus* was stimulated in a dose-dependent manner; it upregulates genes for fermentative acetate, butyrate, and propionate production, as well as group A1 [FeFe] hydrogenases for H_2_ production (5.3- and 4.1-fold at low and high doses, respectively). In contrast, the uncultured genus RUG563 upregulates ethanol production at the expense of downregulating acetate and propionate fermentation pathways. UBA3282 was stimulated only under low 3-NOP dosage. It upregulates group A1 and C2 [FeFe]-hydrogenases, acetate and butyrate production pathways, and pyruvate formate lyase (*pfl*) instead of pyruvate-ferredoxin oxidoreductase (*por*); *pfl* catalyses the non-redox conversion of pyruvate and CoA to acetyl-CoA and formate, thus bypassing the ferredoxin pool and also potentially contributing to the observed formate accumulation following 3-NOP supplementation. In general, the fermentative taxa that were inhibited following 3-NOP supplementation are those that encode limited fermentation routes, including UBA1066, RUG420, *Limivicinus*, *Scatovivens*, and UBA1367 (**Fig. S4b**). However, exceptions include the relatively flexible lineages *Limimorpha* and *Butyrivibrio*_A that were overall suppressed despite each being capable of producing multiple SCFAs, though the latter notably only encodes a single group B [FeFe]-hydrogenase (**Fig. S4b**). We note that fermentative protists and fungi also likely contribute to H_2_ production in these calves, though their abundance and expression was low (<1.2% and <1.1% of metagenomic and metatranscriptomic reads, respectively) and not significantly affected by 3-NOP supplementation.

### Methanogenesis inhibition stimulates growth of novel uncultivated acetogens

Given the short-read analysis (**Fig. 2d, f**) and pathway-centric analysis (**Fig. 4a**) suggest that reductive acetogenesis was stimulated by 3-NOP supplementation, we investigated which species were responsible. To do so, we identified potential hydrogenotrophic acetogen species based on them encoding *cooS* and *acsB*, structural subunits of the signature carbon monoxide dehydrogenase / acetyl-CoA synthase enzyme complex mediating acetogenesis. These species span 26 genera and seven families and were predominantly classified as Clostridia (**Fig. S5**). These acetogens are mainly represented by uncultivated genera not known for mediating acetogenesis, except for *Blautia*^60^. Phylogenetic analysis showed that the AcsB and CooS proteins from these bacteria are canonical but form novel clusters distinct from cultured acetogens such as *Acetobacterium*, *Thermoanaerobacter*, *Clostridium*, and *Moorella* (**Fig. 5d, S6**).

**Figure 5:**
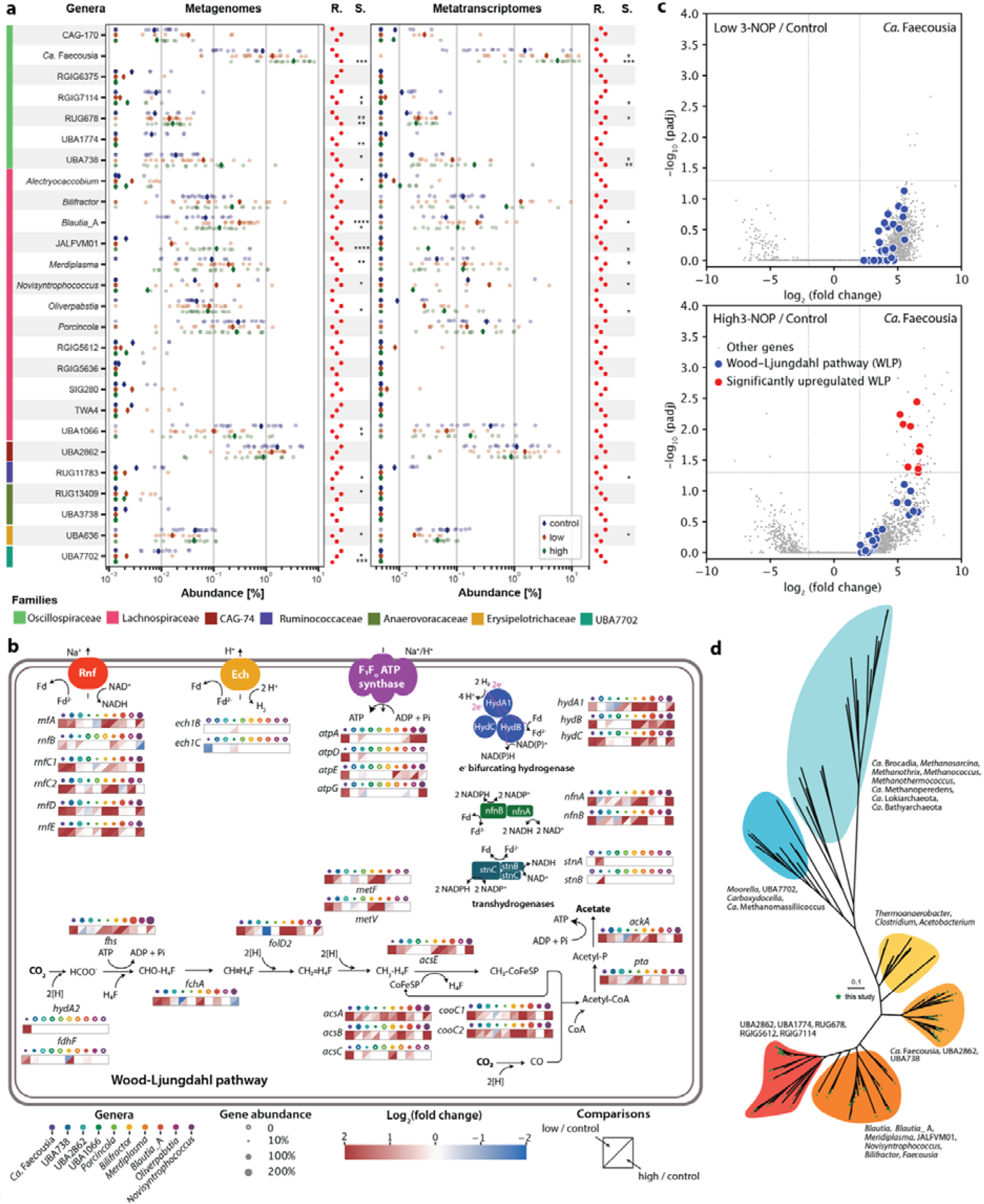
Effects of 3-NOP on the mediators and enzymes of acetogenesis. (**a**) Scatter plot depicting the abundance and activity of acetogen taxa summarised at the genus level, based on mapping against metagenomes and metatranscriptomes. The mean abundance values are indicated by diamonds, while a red dot plot illustrates rank abundance from low (left) to high (right). The statistical significance of differences between groups (low versus control and high versus control) for MAGs was assessed using the Mann-Whitney U test (*p < 0.05, **p < 0.01, ***p < 0.001, ****p < 0.0001). (**b**) A metabolic diagram illustrating the presence of enzymes involved in acetogenesis within the top 10 most abundant acetogen genera, selected based on their mean relative abundance across all samples. The abundance of each gene was calculated as copy number percent aggregated by the genus-level taxonomy of the microorganisms encoding these genes, differentiated by colour. The heatmap indicates the expression fold change (low / control at top left corner and high / control at the bottom right corner) for specific genes, which was averaged at the genus level. CH≡H_4_F, methenyl-tetrahydrofolate; CH_2_=H_4_F, methylene-tetrahydrofolate; CH_3_-H_4_F, methyl-tetrahydrofolate; CHO-H_4_F, formyl-tetrahydrofolate; CoFeSP, corrinoid iron–sulphur protein. (**c**) Volcano plots of the ratio of transcripts for *Ca*. Faecousia under low 3-NOP versus control (top) and high 3-NOP versus control (bottom). Each gene is represented by a grey dot; genes from the Wood-Ljungdahl pathway (WLP) are indicated by blue dots if their differential expression is insignificant (adjusted p ≥ 0.05), or by red dots when significant. The horizontal dotted line indicates an adjusted p value of 0.05, and the vertical dotted line indicates two-fold changes. (**d**) An unrooted phylogenetic tree consisting of 113 AcsB (acetyl-CoA synthase) proteins encoded by the acetogens MAGs identified in this study (indicated by green stars) along with 60 reference sequences from diverse microorganisms, maximising the phylogenetic coverage of AcsB. The tree was inferred using the maximum likelihood method (LG+I+G4) with 1,000 bootstrap iterations after selecting the best substitution model.

Consistent with short-read analysis, putative hydrogenotrophic acetogen species were the most significantly increased microorganisms when methanogens were inhibited by 3-NOP. Putative hydrogenotrophic acetogens accounted for 1.5%, 2.9%, and 4.2% of microbes based on metagenomic read mapping and 1.8%, 6.1%, and 6.5% based on metatranscriptomic read mapping at no, low, and high 3-NOP, respectively. The most abundant and transcriptionally active genera include *Candidatus* (*Ca.*) Faecousia, *Blautia*_A, UBA1066 (Lachnospiraceae), and UBA2862 (Christensenellaceae). Variable responses were observed at the genus level: *Ca.* Faecousia significantly increased in abundance and expression in a dose-dependent manner, with strong stimulation also observed for *Blautia*_A, *Merdiplasma*, JALFVM01 (Lachnospiraceae), and UBA738 (Oscillospiraceae), whereas other lineages were not significantly affected and certain lineages such as UBA1066 decreased in abundance and expression (**Fig. 5a**). This suggests that inhibiting methanogenesis has complex effects on the overall niches for acetogens. However, some taxa are able to strongly respond to H_2_ accumulation and collectively enable acetogens to become the most abundant known hydrogenotrophs in the rumen.

Metabolic annotation of these acetogens shows that the WLP and associated energy conservation machinery is well conserved, including the electron-bifurcating hydrogenase, ferredoxin:NAD-oxidoreductase (Rnf) complex, and F_o_F_1_ ATP synthase, and predominantly upregulated upon 3-NOP supplementation with taxa-specific variations (**Fig. 5b**). Multiple acetogens upregulated the electron-bifurcating group A3 [FeFe]-hydrogenase, which extracts energy from the oxidation of H_2_ (operating in the opposite direction to the equivalent enzyme in fermenters), generating NAD(P)H and reduced ferredoxin to support acetogenesis^61–65^. This upregulation was primarily observed in *Ca.* Faecousia, *Bilifractor*, *Merdiplasma*, and *Novisyntrophococcus*. Additionally, *Ca.* Faecousia, UBA738, *Bilifractor*, *Merdiplasma* and *Blautia*_A upregulated the Rnf complex to mediate electron transfer from reduced ferredoxin, creating a transmembrane sodium or proton concentration gradient to drive ATP synthesis via the ATP synthase^65^. The energy-converting hydrogenase (Ech) hydrogenase complex for energy conservation^65^ was present only in two acetogenic genera and upregulated in only UBA1066 and only under high 3-NOP dosages. The hydrogen-dependent carbon dioxide reductase (HDCR), a filamentous complex mediating hydrogen-dependent CO_2_ reduction to formate^66–68^, was only encoded by *Ca*. Faecousia among rumen acetogens. The acetogens lacking HDCR, including UBA738, *Bilifractor*, and *Blautia*_A, are likely either capable of mixotrophic growth, generating formate through the pyruvate formate lyase reaction, or metabolising formate through metabolic cross-feeding in the rumen. Overall, we observed dose-dependent upregulation of the WLP in *Ca.* Faecousia based on differential expression analysis (**Fig. 5c**).

### Acetogenesis stimulation is a common response to methanogenesis inhibition

To determine if these findings extend to other ruminants, we reanalysed a previously reported field trial on 3-NOP focused on Holstein cows, where methane emissions were also reduced^28^. In line with the present study, analysis of metagenomic short reads confirmed that the abundance and expression of MCR, methanogenic hydrogenases, and electron-bifurcating hydrogenases decreased. In parallel, we observed an increase in the abundance and expression of the acetyl-CoA synthase / carbon monoxide dehydrogenase complex for acetogenesis, together with fermentative and sensory [FeFe]-hydrogenases (**Fig. S7a**). In line with the differential efficacy of 3-NOP, the differences in gene and transcript abundance were more modest for the Holstein study (21% reduction in methane yield) than the present Holstein X Jersey study (62% reduction). Mapping of the genome catalogue to metagenomic and metatranscriptomic reads confirmed a range of acetogens were enriched following 3-NOP supplementation in the Holstein study, most significantly *Porcinola*, but also UBA2862, RUG678, and RUG13409 (**Fig. S7b**). However, we did not detect reads from *Ca*. Faecousia, which was the most abundant acetogen in the current Holstein X Jersey study. Collectively, these results suggest 3-NOP broadly inhibits methanogenesis, remodels fermentation, and stimulates acetogenesis, though the taxa involved vary between studies. It is unclear whether other methanogenesis inhibitors also stimulate acetogenesis, with a recent study suggesting that respiratory rather than acetogenic hydrogenotrophs are stimulated in the rumen upon administration of *Asparagopsis*, containing the methanogenesis inhibitor bromoform^10^.

## Conclusions

Here we provide the most detailed species-resolved view to date of the response of rumen microbiota and processes to methanogenesis inhibition. Our data shows that microbiota remodelling explains why 3-NOP, despite being a potent methane inhibitor, maintains rather than enhances host productivity. 3-NOP potently and specifically inhibits diverse methanogens *in situ*, leading to the accumulation of their primary substrates such as H_2_ and formate. Hydrogenotrophic and fermentative bacteria alike respond to H_2_ accumulation through both community shifts and modulating gene expression. The most stimulated microbes are diverse uncultivated lineages of Clostridia that mediate reductive acetogenesis via the WLP. On the other hand, the accumulation of H_2_ causes fermenters to remodel their pathways away from acetate production towards undesirable compounds such as ethanol. In addition to the direct loss of energy as H_2_ and CH_4_ (approximately 1.5% and 2.5% of feed energy content, respectively), the divergent responses of H_2_-cycling microbes mean the overall SCFA pool doesn’t significantly increase for ruminants to absorb energy from. This provides a biological explanation for why methanogenesis inhibition leads to negligible productivity gains.

Optimising methanogenesis inhibition, such that electrons flow into absorbable nutrients rather than methane, can be facilitated by microbiota-wide approaches that consider all community members to the direct and indirect effects of methanogenesis inhibitors. In this regard, it is critical to consider uncultured bacteria, given our study emphasises they are dominant but overlooked fermenters and hydrogenotrophs in the rumen, including expanding knowledge of the diversity and physiology of gastrointestinal acetogens beyond culture-based studies^4,43^. Genome-informed cultivation efforts are essential to better understand the physiology of these microbes and determine how we can leverage them. From our perspective, there are at least two promising solutions to redirect electron flow in the rumen to enhance productivity following methanogenesis inhibition. First, stimulating respiratory microbes, for example *via* addition of limiting electron acceptors, to reduce H_2_ levels and emissions such that fermentation occurs through the most efficient route. Second, inhibiting metabolically flexible fermenters through targeting their regulatory hydrogenases, to promote fermentative SCFA formation and reduce H_2_ build up that creates the negative feedback loops observed in this study. The analytical framework and extensive MAG collection provided here are valuable resources to guide these efforts.

## Materials and Methods

### Animal field trials, ruminal H_2_ and CH_4_ measurements, short-chain fatty acid assays and performance data

A trial with three groups was conducted at AgResearch in Palmerston North, New Zealand. Fifty-one animals (Holstein X Jersey female calves) were enrolled at birth, and randomly assigned to three treatment groups based on date of birth, birth weight, and parity of the mother, resulting in three homogeneous groups of 17 animals each: Group 1: Control, No 3-NOP; Group 2: Low 3-NOP (target dose 150 mg 3-NOP / kg dry matter intake (DMI) per day; Group 3: High 3-NOP (target dose 250 mg 3-NOP / kg DMI per day). 3-NOP was added in both the calf milk replacer (CMR) and also the calf starter feed (CS) and both feed sources were offered daily on an ad lib basis. In addition, all animals had access to hay which was not measured as it was supplied on a group basis and was not supplemented with any of the test items, but was estimated to make up approximately 7% of the total diet. Individual feed intake of calf milk replacer and calf starter was measured daily for the duration of the study, including during the period in the respiration chamber. Animals remained in study for a total of 103 ± 3 days and were weaned (i.e., no longer received CMR) after 82 ± 3 days of age. Animals were placed into respiration chambers when they were 97 days of age on average, and gas emissions (CH_4_, H_2_, and CO_2_) were measured over a 48-h period at the end of the study. When animals were removed from the chambers at 99 ± 1 days of age, they were weighed and rumen samples were removed via oesophageal tubing and split into different aliquots to be used either for determination of SCFAs, other rumen fermentation metabolites, and for DNA and RNA extraction. All procedures involving animals were conducted in accordance with the established ethical guidelines of the trial site. Animal phenotypic data is presented in **Dataset S6**.

### Library preparation and sequencing

Paired-end metagenomic reads from 51 calves were collected at AgResearch in New Zealand, with 44 of these samples also providing matching metatranscriptomic reads. Rumen samples were collected from all calves at approximately 99 ± 1 days of age via oesophageal tubing. Subsamples for short-chain fatty acid (SCFA), DNA, and RNA analysis were collected. RNA samples were snap-frozen in liquid nitrogen and immediately transferred to a −80°C freezer for storage until analysis. Other samples were immediately placed on ice after collection; DNA subsamples were frozen at -20°C, while SCFA samples were processed without delay. Both metagenomic and metatranscriptomic samples were processed according to the methods described by Shi et al (2014)^69^. Community DNA and RNA were purified from ruminal samples of cannulated calves following established protocols^17^. For metagenomic sequencing, DNA was prepared using the Nextera DNA Library Prep Kit (Illumina) and sequenced on an Illumina platform. For metatranscriptomics, ribosomal RNA was depleted from total RNA with the Ribo-Zero Plus rRNA Removal Kit (Illumina). Double-stranded cDNA was synthesised from mRNA-enriched RNA using the TruSeq Stranded mRNA Kit (Illumina), then a library was constructed and sequenced on an Illumina platform. Raw metagenomic reads underwent quality control using BBDuk (with BBMap v39.00)^70^, which involved trimming adapters, filtering PhiX reads, trimming the 3’ ends, setting a Phred quality threshold of 15 (96.8% base call accuracy), and discarding reads shorter than 50 bp in length. Read quality assessment was performed using FastQC (v0.11.9)^71^ coupled with MultiQC (v1.13)^72^. Summary statistics for metagenomic and metatranscriptomic reads, both pre- and post-quality control, are detailed in **Dataset S7**.

### Metagenomic assembly and binning

Assembly and binning were performed by using the quality filtered reads from the 51 metagenomes. To maximise the representation of ruminal microbial diversity, two additional sequencing datasets were included (**Dataset S8**). The first dataset, an unpublished collection of 32 metagenomic libraries, was from eight calves subjected to 3-NOP supplementation and various diets. The second dataset, from Pitta *et al*., (2022)^17^, comprises 24 matching metagenomic and metatranscriptomic libraries from eight calves subjected to either control or 3-NOP dosages. The sequencing reads from this dataset were retrieved from the NCBI Sequence Read Archive (SRA) under accession PRJNA666417. For each dataset, processed reads were individually assembled and binned for each library. Metagenomic assembly was carried out using metaSPAdes (v3.15.5)^73^ with a custom set of k-mers, ‘-k 27,37,47,57,67,77,87,97,107,117,127’. This was followed by binning with MetaBAT2 (v2.12.1)^74^, MaxBin2 (v2.2.7)^75^, and CONCOCT (v1.1.0)^76^ to reconstruct metagenome-assembled genomes (MAGs), which are implemented in the ‘binning’ module of MetaWRAP (v1.3.2)^77^. Subsequently, the bin refinement module of MetaWRAP was used to consolidate the MAGs from each assembly into a medium-to-high quality set with a minimum completion of 80% and a maximum contamination of 10%. A dereplicated set of MAGs was produced in each study using dRep’s ‘dereplicate’ function (v3.4.2)^78^. Completeness and contamination of the MAGs were estimated using CheckM2 ‘predict’ v0.1.3^79^. Taxonomy assignment for each MAG was performed using GTDB-Tk ‘classify_wf’ (v2.1.1)^80^. To maximise representation of ruminal microbiota, a consolidated set of genomes was created, comprising metagenome-assembled genomes (MAGs) from the present study and publicly available rumen microbial pure culture genomes and MAGs^43–50^. These MAGs were mapped to metagenomes and metatranscriptomes to assess the abundance and activity of key microbes mediating digestive processes in the rumen and were linked to ruminant GHG emissions and productivities. Genome dereplication was performed using dRep, with parameters ‘-comp 50 -con 10 -sa 0.95 -nc 0.3’ to delineate species-level bins at 50% completion and 10% contamination thresholds, which also recognises MAGs that have lower levels according to Almeida *et al*., (2019)^81^ and Watson (2021)^82^. Subsequently, the CoverM ‘genome’ function was used to calculate the relative abundance of each MAGs in each sample^83^. CoverM ‘genome’ was also used to estimate the activity of each organism in each sample by mapping high-quality metatranscriptomic reads to the MAGs. Full details of the MAGs, including genome taxonomy, quality, metabolic annotation, relative abundance measurements, functional group assignments, and species placeholder names, are provided in **Dataset S2**.

### Microbial community composition and diversity analysis

Microbial community composition was analysed using SingleM (version 1.0.0beta2)^84^, which incorporates 59 single-copy ribosomal marker genes. The ribosomal protein gene *rpsB*, which is conserved across bacteria and archaea, was selected to generate a table of gene counts for each operational taxonomic unit (OTU table) for subsequent analyses. Community alpha and beta diversity metrics were calculated using the VEGAN^85^ package in RStudio. The OTU table was rarefied to match the minimum sequence count observed across samples. For alpha diversity, observed and estimated richness (Chao1) were computed for each sample, with differences across groups assessed using a non-parametric statistical test. For beta diversity, community structure distances were quantified using the Bray-Curtis dissimilarity method and visualised with multidimensional scaling ordination techniques. Differences in community structure between groups were evaluated for significance using permutational multivariate analysis of variance (PERMANOVA). Fungal population abundance was estimated by determining the proportion of short reads aligning to a set of single-copy ribosomal proteins, facilitated by SingleM^84^, in conjunction with PhyloFlash^86^, which searches the SILVA ribosomal RNA small subunit (SSU) database (v138)^87^. Eukaryotic taxonomy across three metagenomes was determined using PhyloFlash, which queried gut fungal and protist databases.

### Metabolic annotation of short reads

Metabolic capabilities were inferred from metagenomic and metatranscriptomic datasets by annotating high-quality short reads using DIAMOND (-max-target-seqs 1, -max-hsps 1)^88^ aligned against an in-house database of 51 metabolic marker proteins with carefully-curated filtering thresholds^6,89–91^, as well as against the KEGG database. Ribosomal RNA was removed from high-quality metatranscriptomic short reads using RiboDetector (v0.2.7)^92^, followed by the removal of reads aligning to the Jersey cattle reference genome (accession GCA_021234555.1) using BBDuk^70^. Gene abundance from short reads was calculated as an indicator of metabolic capabilities at a community level. It was quantified as ‘average gene copies per organism’ by dividing the abundance of the gene (in reads per kilobase million, RPKM) by the mean abundance of 14 universal single-copy ribosomal marker genes (in RPKM, obtained from the SingleM v0.13.2 package)^84^. Sequencing depth was calculated using Seqkit’s ‘stat’ function (v2.3.0)^93^. Non-parametric Kruskal-Wallis tests were used to determine statistical significance across groups, adjusting *p*-values with the False Discovery Rate method via the Benjamini-Hochberg procedure, implemented in the R ’stats’ and ’broom’ packages^94^. Data processing and visualisation were conducted using the ‘tidyverse’^95^ and ‘ggplot2’^96^ packages in R (v4.2.3).

### Metabolic annotation of MAGs

For the consolidated MAG set generated in this study, in-depth metabolic annotation was conducted using predicted proteins from each MAG. Firstly, an in-house approach for reductive acetogenesis via the Wood-Ljungdahl pathway (WLP) was determined by screening for marker genes for acetyl-CoA synthase (*acsB*) and anaerobic carbon monoxide dehydrogenase (*cooS*). Reference sequences from diverse lineages^90^, along with all genes associated with the WLP and energy conservation in *Acetobacterium woodii*^97^*, Thermoanaerobacter kivui*^98^, and *Sporomusa ovata*^99^ served as references. A quality control cut-off of 80% alignment length was applied. Secondly, all KO terms for both reductive and oxidative branches of acetate formation at the Kyoto Encyclopedia of Genes and Genomes (KEGG)^100^ were identified (**Dataset S5)**. The high level of agreement in the results of reductive acetogenesis between the two methods demonstrates the accuracy of this approach. Fermentative capabilities for butyrate and propionate (via acrylate or succinate) were subsequently established using the associated KO terms, complemented with the identification of group A1, A3, or B [FeFe] hydrogenases from the MAGs. KO terms were annotated from the MAGs using DRAM. Methanogens were identified as archaea encoding McrA. The statistical significance of relative abundance across groups, at both metagenomic and metatranscriptomic levels, was tested using the Mann–Whitney U test, implemented in custom Python (v3.10.10) scripts with the ’numpy’^101^, ’scipy’^102^, and ’statsmodels’^103^ packages. The distribution and mean abundance values of each taxon across groups were visualised using scatter plot functions from the ‘matplotlib’^104^ and ‘seaborn’^105^ libraries in Python. To facilitate log transformations during visualisation, a pseudo-zero, defined as half of the minimal value of all relative abundance values in each analysis, was applied.

### Metatranscriptomic expression analysis

An overall mapping rate of 65.7% was achieved based on predicted gene sequences from MAGs using Salmon (v 1.10.2)^106^. Differential expression analysis of transcripts was carried out using pydeseq2 (v 0.4.9)^107^ to calculate fold change and adjusted *p*-values. Transcripts were categorised into SCFA production pathways using their corresponding KO terms as summarised in **Dataset S5**. The log2 transformed fold change values between high 3-NOP vs control and low 3-NOP vs control for fermentation genes and hydrogenases are summarised in **Dataset S4**.

### Phylogenetic tree inferences

These annotations were complemented by evolutionary analyses employing both phylogenetic (gene-based) and phylogenomic (MAG-based) methodologies. Phylogenetic trees were constructed for the methyl-CoM reductase (McrA), AcsB, CooS, and various hydrogenases ([NiFe], [Fe], and [FeFe]) using protein sequences obtained from MAGs through homology-based screening against our proprietary database of metabolic marker genes. Alignment of protein sequences was performed using MAFFT with reference sequences, followed by trimming using trimAl ’gappyout’ (v1.4.rev15)^108^. The resulting trimmed multiple sequence alignments were utilized for phylogenetic inference in IQ-TREE with parameters ’alrt 1000 -B 1000 -m TEST’ (v2.2.0.3)^109^, including evolutionary model testing and 1000 bootstrap iterations. Visualisation of the resulting trees was accomplished using iTOL (v6)^110^, complemented by custom Python scripts for tree annotation and styling. Phylogenomic trees for methanogens and acetogens were inferred from their respective MAGs using PhyloPhlAn^111,112^, which employs 400 universal proteins for multiple sequence alignment on MAFFT ‘--anysymbol’ (v7.508)^113,114^. This was followed by alignment trimming with trimAl and tree inference with IQ-TREE, employing the parameters previously described. Visualisation and annotation of phylogenomic trees were conducted using iTOL, with final rendering conducted using custom Python scripts with ‘matplotlib’^104^ and ‘seaborn’^105^ packages.

### Docking simulations of 3-NOP in AlphaFold2 generated models of MCR

We examined all protein sequences for the subunits of MCR predicted from MAGs, retaining 64 complete sequences for structure prediction using FastFold (v0.2.0)^115^, with weights from AlphaFold (v2.3)^116^ incorporated as individual monomers. Hexameric complexes, including cofactors, were constructed by superposing each subunit onto a crystallographic template from the Protein Data Bank (PDB ID: 1HBO) via the structural alignment script in the Maestro Small-Molecule Drug Discovery Suite (v2021-2). Initially, we compared this procedure with the computationally more intensive one-shot prediction of hexamers by AlphaFold Multimer (v3)^116^. Given that the root mean-square deviation (RMSD) between the two methods was only 0.48 Å for the protein backbone atoms, we opted to continue with the superposition procedure. Subsequently, hydrogen atoms were added using the Protein Preparation Wizard in Maestro^117^, with heteroatom protonation set at a pH of 6.7, matching the intracellular pH of methanogenic archaea^118^. Finally, the hydrogen bonding network was reoriented using the PROPKA routine at pH 6.7, and the structures underwent restrained minimisation with a threshold of 0.3 Å for protein heavy atoms, employing the OPLS4e force field. Docking of 3-NOP was conducted using Glide in standard precision mode^119^, with the coenzyme-B (CoB) cofactor retained within the structures. The search space covered the complete active site of the enzyme, with a centroid close to the sulfur atom of CoB. The accuracy of the docking protocol was validated through cross-docking with the coenzyme-M cofactor and assessed using the rmsd.py routine included with Maestro, as recommended^120^. Validation indicated excellent pose prediction accuracy (RMSD < 1.0 Å), with a valid pose of CoM found in all 64 structures. Accordingly, poses of 3-NOP were classified into forward and reversed binding modes based on an RMSD threshold of 1.0 Å to docking poses obtained from our previous work. For visualisation, only complexes with a highly accurate 3-NOP binding orientation, placing the nitrate ester moiety closely to the nickel ion in the F_430_ cofactor, were selected.

### Calculations of combustion energy, reductant recovery and energy loss

Combustion energy per litre of rumen fluid were calculated by multiplying the concentration of each SCFA with the Gibbs free energy released from their aerobic oxidation at standard conditions (ΔG^0^’)^25^ according to equations (**Eq. 1-5**) as follows:

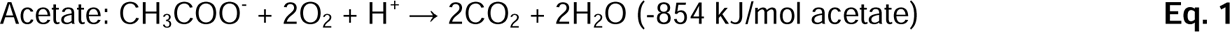

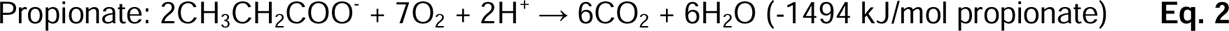

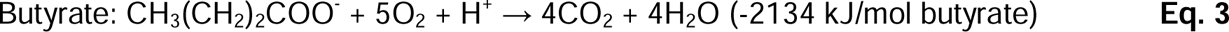

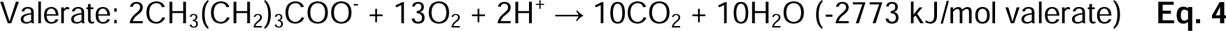

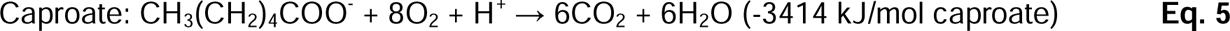

Amounts of reductant (i.e., the amounts of reducing equivalents that become available upon complete oxidation of a compound) for CH_4_ and H_2_ emitted by calves per day were calculated by dividing daily emitted masses by the molecular weight of the respective gas (16 g/mol for CH_4_ and 2 g/mol for H_2_) multiplied by 8 and 2 for CH_4_ and H_2_, respectively. Subsequently, mean amounts of reductant emitted as H_2_ by calves of the control group were subtracted from amounts of reductant emitted as H_2_ by individual calves of the low and high 3-NOP treatment groups. These net amounts of H_2_-reductant were divided by the mean amounts of reductant emitted as CH_4_ in the control group. The resulting percentage (i.e., reductant recovered as H_2_) reflects the fraction of reductant that is lost due to enhanced H_2_ emission, the result of partial methanogenesis inhibition, in 3-NOP treated calves relative to the amount of reductant that is lost due to methanogenesis in untreated calves. In addition, reductant recovered as CH_4_ was calculated by dividing amounts of reductant emitted as CH_4_ in individual calves of 3-NOP treatment groups with mean amounts of reductant emitted as CH_4_ in the control group. Energy lost by the emission of H_2_ and CH_4_ were calculated based on the Gibbs free energy (ΔG^0^’)^25^ released during aerobic oxidation of H_2_ (**Eq. 6**) and CH_4_ and CH_4_ (**Eq. 7**), respectively.

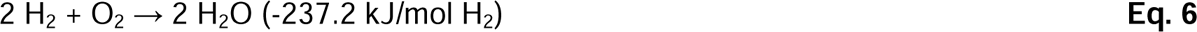

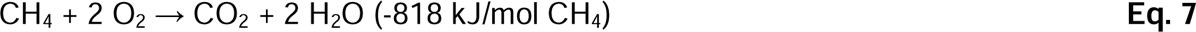

## Footnotes

## Conflict of interest statement

This study was primarily funded by dsm-firmenich. C.G., R.G. and G.N. receive funding from dsm-firmenich. N.W., M.K., E.V.L., M.W., A.F., and R.S. are employees of dsm-firmenich.

## Author contributions

C.G., N.W., M.K., E.V.L., and G.N. designed this study. N.W., M.K., and E.V.L. coordinated animal trials, phenotypic analysis, and meta-omic sequencing. G.N., C.G., and M.W. developed MAG collection. G.N. and C.G. conducted metabolic annotations and phylogenetic analysis with input from S.J., O.S., and M.J. O.S. and R.K.T. contributed energy calculations. A.F., R.G., R.S., and G.N. conducted docking analysis. V.M., M.W., R.K.T., and P.B.P. contributed datasets and insights for metabolic analyses. G.N. and C.G. wrote the manuscript with input from all authors.

## Data and code availability

Raw sequencing reads are deposited in the NCBI Sequence Read Archive (SRA) under Bioproject accession. Gas chamber measurements, phenotypic profiling, animal performance, and meta-Omics analysis scripts are publicly available on GitHub.

## Acknowledgements

This study was supported by dsm-firmenich, an NHMRC EL2 Fellowship (APP1178715; awarded to C.G) and a Monash Faculty of Medicine, Nursing, and Health Sciences Early Career Postdoctoral Fellowship 2023 (ECPF23–8566329039; awarded to G.N.). We gratefully acknowledge the support of Carlos Ramirez-Palacios in setting up the FastFold and AlphaFold multimer workflows and providing the structure files. We are grateful for helpful discussions with Rudolf Thauer, Tom Watts, Caitlin Welsh, Bob Leung, and Francesco Ricci.

## References

1. Ruminant Methanogens as a Climate Change Target. ASM.org https://asm.org:443/Articles/2023/June/Ruminant-Methanogens-as-a-Climate-Change-Target.

2. United Nations Environment Programme/Climate and Clean Air Coalition. Global Methane Assessment: 2030 Baseline Report. https://wedocs.unep.org/bitstream/handle/20.500.11822/41107/methane_2030.pdf?sequence=1&isAllowed=y (2022).

3. Johnson, K. A. & Johnson, D. E. Methane Emissions from Cattle. Journal of Animal Science 73, 2483–2492 (1995).

4. Mizrahi, I., Wallace, R. J. & Moraïs, S. The rumen microbiome: balancing food security and environmental impacts. Nature Reviews Microbiology 19, 553–566 (2021).

5. Dijkstra, J. Production and absorption of volatile fatty acids in the rumen. Livestock Production Science 39, 61–69 (1994).

6. Greening, C. et al. Diverse Hydrogen Production and Consumption Pathways Influence Methane Production in Ruminants. The ISME Journal 13, 2617–2632 (2019).

7. Stams, A. J. M. & Plugge, C. M. Electron transfer in syntrophic communities of anaerobic bacteria and archaea. Nature Reviews Microbiology 7, 568 (2009).

8. Kelly, W. J. et al. Hydrogen and Formate Production and Utilisation in the Rumen and the Human Colon. Animal Microbiome 4, 22 (2022).

9. Cord-Ruwisch, R., Seitz, H.-J. & Conrad, R. The capacity of hydrogenotrophic anaerobic bacteria to compete for traces of hydrogen depends on the redox potential of the terminal electron acceptor. Archives of Microbiology 149, 350–357 (1988).

10. Zhang, P. et al. Red seaweed supplementation suppresses methanogenesis in the rumen, revealing potentially advantageous traits among hydrogenotrophic bacteria. Preprint at 10.1101/2024.06.07.597961 (2024).

11. Hristov, A. N. et al. An inhibitor persistently decreased enteric methane emission from dairy cows with no negative effect on milk production. Proceedings of the National Academy of Sciences 112, 10663–10668 (2015).

12. Duin, E. C. et al. Mode of action uncovered for the specific reduction of methane emissions from ruminants by the small molecule 3-nitrooxypropanol. Proceedings of the National Academy of Sciences 113, 6172–6177 (2016).

13. Hristov, A., et al. Mitigation of Greenhouse gas Emissions in Livestock Production: A Review of Technical Options for Non-CO2 Emissions; FAO Anim. Produon and Health Paper 1–206.

14. Vyas, D. et al. The Combined Effects of Supplementing Monensin and 3-nitrooxypropanol on Methane emissions, Growth rate, and Feed Conversion Efficiency in Beef Cattle Fed High-forage and High-grain diets. Journal of Animal Science 96, 2923–2938 (2018).

15. Alemu, A. W. et al. 3-Nitrooxypropanol Decreased Enteric Methane Production From Growing Beef Cattle in a Commercial Feedlot: Implications for Sustainable Beef Cattle Production. Frontiers in Animal Science 2, (2021).

16. Almeida, A. K. et al. Effect of 3-nitrooxypropanol on enteric methane emissions of feedlot cattle fed with a tempered barley-based diet with canola oil. Journal of Animal Science 101, skad237 (2023).

17. Pitta, D. W. et al. The effect of 3-nitrooxypropanol, a potent methane inhibitor, on ruminal microbial gene expression profiles in dairy cows. Microbiome 10, 146 (2022).

18. Van Wesemael, D. et al. Reducing enteric methane emissions from dairy cattle: Two ways to supplement 3-nitrooxypropanol. Journal of Dairy Science 102, 1780–1787 (2019).

19. Haisan, J. et al. The effects of feeding 3-nitrooxypropanol at two doses on milk production, rumen fermentation, plasma metabolites, nutrient digestibility, and methane emissions in lactating Holstein cows. Anim. Prod. Sci. 57, 282–289 (2016).

20. Melgar, A. et al. Short communication: Short-term effect of 3-nitrooxypropanol on feed dry matter intake in lactating dairy cows. Journal of Dairy Science 103, 11496–11502 (2020).

21. Martinez-Fernandez, G. et al. 3-NOP vs. Halogenated Compound: Methane Production, Ruminal Fermentation and Microbial Community Response in Forage Fed Cattle. Frontiers in Microbiology 9, (2018).

22. van Gastelen, S. et al. 3-Nitrooxypropanol decreases methane emissions and increases hydrogen emissions of early lactation dairy cows, with associated changes in nutrient digestibility and energy metabolism. Journal of Dairy Science 103, 8074–8093 (2020).

23. van Gastelen, S. et al. Long-term effects of 3-nitrooxypropanol on methane emission and milk production characteristics in Holstein Friesian dairy cows. Journal of Dairy Science (2024) doi:10.3168/jds.2023-24198.

24. Melgar, A. et al. Dose-response effect of 3-nitrooxypropanol on enteric methane emissions in dairy cows. Journal of Dairy Science 103, 6145–6156 (2020).

25. Thauer, R. K., Jungermann, K. & Decker, K. Energy conservation in chemotrophic anaerobic bacteria. Bacteriol Rev 41, 100–180 (1977).

26. Meale, S. J. et al. Early life dietary intervention in dairy calves results in a long-term reduction in methane emissions. Scientific Reports 11, 3003 (2021).

27. Furman, O. et al. Stochasticity constrained by deterministic effects of diet and age drive rumen microbiome assembly dynamics. Nature Communications 11, 1904 (2020).

28. Melgar, A. et al. Effects of 3-nitrooxypropanol on rumen fermentation, lactational performance, and resumption of ovarian cyclicity in dairy cows. Journal of Dairy Science 103, 410–432 (2020).

29. Zhang, X. M. et al. 3-Nitrooxypropanol supplementation had little effect on fiber degradation and microbial colonization of forage particles when evaluated using the in situ ruminal incubation technique. Journal of Dairy Science 103, 8986–8997 (2020).

30. Ungerfeld, E. M. Shifts in metabolic hydrogen sinks in the methanogenesis-inhibited ruminal fermentation: a meta-analysis. Frontiers in Microbiology 6, (2015).

31. Rea, S., Bowman, J. P., Popovski, S., Pimm, C. & Wright, A.-D. G. *Methanobrevibacter millerae* sp. nov. and *Methanobrevibacter olleyae* sp. nov., Methanogens From the Ovine and Bovine Rumen That Can Utilize Formate For Growth. International Journal of Systematic and Evolutionary Microbiology 57, 450–456 (2007).

32. Kelly, W. J. et al. Occurrence and Expression of Genes Encoding Methyl-compound Production in Rumen Bacteria. Animal Microbiome 1, 15 (2019).

33. Veresegyházy, T., Fébel, H., Nagy, G. & Rimanóczy, Á. Disappearance of ethanol from isolated sheep rumen. Acta Veterinaria Hungarica 51, 189–196 (2005).

34. Seedorf, H. et al. The genome of Clostridium kluyveri, a strict anaerobe with unique metabolic features. Proceedings of the National Academy of Sciences 105, 2128–2133 (2008).

35. Thauer, R. K., Jungermann, K., Henninger, H., Wenning, J. & Decker, K. The Energy Metabolism of *Clostridium kluyveri*. European Journal of Biochemistry 4, 173–180 (1968).

36. Thauer, R. K. My Lifelong Passion for Biochemistry and Anaerobic Microorganisms. Annual Review of Microbiology 69, 1–30 (2015).

37. Raun, B. M. L. & Kristensen, N. B. Metabolic effects of feeding ethanol or propanol to postpartum transition Holstein cows. Journal of Dairy Science 94, 2566–2580 (2011).

38. Emery, R. S., Lewis, T. R., Everett, J. P. & Lassiter, C. A. Effect of Ethanol on Rumen Fermentation1. Journal of Dairy Science 42, 1182–1186 (1959).

39. Gruninger, R. J. et al. Application of 3-nitrooxypropanol and canola oil to mitigate enteric methane emissions of beef cattle results in distinctly different effects on the rumen microbial community. anim microbiome **4**, 35 (2022).

40. Guyader, J., Ungerfeld, E. M. & Beauchemin, K. A. Redirection of Metabolic Hydrogen by Inhibiting Methanogenesis in the Rumen Simulation Technique (RUSITEC). Front. Microbiol. 8, (2017).

41. Zijderveld, S. M. V. et al. Nitrate and sulfate: Effective alternative hydrogen sinks for mitigation of ruminal methane production in sheep. Journal of Dairy Science 93, 5856–5866 (2010).

42. Bayaru, E. et al. Effect of fumaric acid on methane production, rumen fermentation and digestibility of cattle fed roughage alone. Nihon Chikusan Gakkaiho 72, 139–146 (2001).

43. Xie, F. et al. An integrated gene catalog and over 10,000 metagenome-assembled genomes from the gastrointestinal microbiome of ruminants. Microbiome 9, 137 (2021).

44. Seshadri, R. et al. Cultivation and sequencing of rumen microbiome members from the Hungate1000 Collection. Nature Biotechnology 36, 359–367 (2018).

45. Peng, X. et al. Genomic and functional analyses of fungal and bacterial consortia that enable lignocellulose breakdown in goat gut microbiomes. Nature Microbiology 6, 499–511 (2021).

46. Mukherjee, S. et al. 1,003 reference genomes of bacterial and archaeal isolates expand coverage of the tree of life. Nature Biotechnology 35, 676–683 (2017).

47. Anderson, C. L. & Fernando, S. C. Insights into rumen microbial biosynthetic gene cluster diversity through genome-resolved metagenomics. Communications Biology 4, 1–12 (2021).

48. Martínez-Álvaro, M. et al. Bovine Host Genome Acts on Rumen Microbiome Function Linked to Methane Emissions. Communications Biology 5, 1–16 (2022).

49. Andersen, T. O. et al. Metabolic influence of core ciliates within the rumen microbiome. ISME J 1–13 (2023) doi:10.1038/s41396-023-01407-y.

50 Cow-rumen catalogue v1.0 released in MGnify. (2021).

51. Li, Q., et al. Reductive acetogenesis is a dominant process in the ruminant hindgut. Preprint at 10.21203/rs.3.rs-4473149/v1 (2024).

52. Thauer, R. K. Methyl (Alkyl)-Coenzyme M Reductases: Nickel F-430-Containing Enzymes Involved in Anaerobic Methane Formation and in Anaerobic Oxidation of Methane or of Short Chain Alkanes. Biochemistry 58, 5198–5220 (2019).

53. Prakash, D., Wu, Y., Suh, S.-J. & Duin, E. C. Elucidating the Process of Activation of Methyl-Coenzyme M Reductase. Journal of Bacteriology 196, 2491–2498 (2014).

54. Angenent, L. T. et al. Chain Elongation with Reactor Microbiomes: Open-Culture Biotechnology To Produce Biochemicals. Environmental Science & Technology 50, 2796– 2810 (2016).

55. J. Steinbusch, K. J. M. Hamelers, H. V. M. Plugge, C. & N. Buisman, C. J. Biological formation of caproate and caprylate from acetateD: fuel and chemical production from low grade biomass. Energy & Environmental Science 4, 216–224 (2011).

56. Spirito, C. M., Richter, H., Rabaey, K., Stams, A. J. & Angenent, L. T. Chain elongation in anaerobic reactor microbiomes to recover resources from waste. Current Opinion in Biotechnology 27, 115–122 (2014).

57. Schut, G. J. & Adams, M. W. W. The Iron-Hydrogenase of *Thermotoga maritima* Utilizes Ferredoxin and NADH Synergistically: a New Perspective on Anaerobic Hydrogen Production. Journal of Bacteriology 191, 4451–4457 (2009).

58. Zheng, Y., Kahnt, J., Kwon, I. H., Mackie, R. I. & Thauer, R. K. Hydrogen Formation and Its Regulation in *Ruminococcus albus*: Involvement of an Electron-Bifurcating [FeFe]-Hydrogenase, of a Non-Electron-Bifurcating [FeFe]-Hydrogenase, and of a Putative Hydrogen-Sensing [FeFe]-Hydrogenase. Journal of Bacteriology 196, 3840–3852 (2014).

59. Pony, P., Rapisarda, C., Terradot, L., Marza, E. & Fronzes, R. Filamentation of The Bacterial Bi-functional Alcohol/Aldehyde Dehydrogenase AdhE is Essential For Substrate Channeling and Enzymatic Regulation. Nature Communications 11, 1426 (2020).

60. Trischler, R., Roth, J., Sorbara, M. T., Schlegel, X. & Müller, V. A Functional Wood– Ljungdahl Pathway Devoid of a Formate Dehydrogenase in The Gut Acetogens *Blautia wexlerae*, *Blautia luti* and Beyond. Environmental Microbiology 24, 3111–3123 (2022).

61. Schuchmann, K. & Müller, V. A Bacterial Electron-bifurcating Hydrogenase *. Journal of Biological Chemistry 287, 31165–31171 (2012).

62. Müller, V., Chowdhury, N. P. & Basen, M. Electron Bifurcation: A Long-Hidden Energy-Coupling Mechanism. Annual Review of Microbiology 72, 331–353 (2018).

63. Katsyv, A. et al. Molecular Basis of the Electron Bifurcation Mechanism in the [FeFe]-Hydrogenase Complex HydABC. Journal of the American Chemical Society 145, 5696–5709 (2023).

64. Buckel, W. & Thauer, R. K. Flavin-Based Electron Bifurcation, A New Mechanism of Biological Energy Coupling. Chemical Reviews 118, 3862–3886 (2018).

65. Schuchmann, K. & Müller, V. Autotrophy at the thermodynamic limit of life: a model for energy conservation in acetogenic bacteria. Nature Reviews Microbiology 12, 809–821 (2014).

66. Schuchmann, K. & Müller, V. Direct and Reversible Hydrogenation of CO_2_ to Formate by a Bacterial Carbon Dioxide Reductase. Science 342, 1382–1385 (2013).

67. Dietrich, H. M. et al. Membrane-Anchored HDCR Nanowires Drive Hydrogen-Powered CO_2_ Fixation. Nature 607, 823–830 (2022).

68. Müller, V. New Horizons in Acetogenic Conversion of One-Carbon Substrates and Biological Hydrogen Storage. Trends in Biotechnology 37, 1344–1354 (2019).

69. Shi, W. et al. Methane yield phenotypes linked to differential gene expression in the sheep rumen microbiome. Genome Research 24, 1517–1525 (2014).

70 Bushnell, B. BBMap. Joint Genome Institute (2022).

71. Andrews, S. FastQC A Quality Control tool for High Throughput Sequence Data. Babraham Bioinformatics.

72. Ewels, P., Magnusson, M., Lundin, S. & Käller, M. MultiQC: summarize analysis results for multiple tools and samples in a single report. Bioinformatics 32, 3047–3048 (2016).

73. Nurk, S., Meleshko, D., Korobeynikov, A. & Pevzner, P. A. metaSPAdes: a new versatile metagenomic assembler. Genome Research 27, 824–834 (2017).

74. Kang, D. D. et al. MetaBAT 2: an adaptive binning algorithm for robust and efficient genome reconstruction from metagenome assemblies. PeerJ 7, e7359 (2019).

75. Wu, Y.-W., Simmons, B. A. & Singer, S. W. MaxBin 2.0: an automated binning algorithm to recover genomes from multiple metagenomic datasets. Bioinformatics 32, 605– 607 (2015).

76. Alneberg, J. et al. Binning metagenomic contigs by coverage and composition. Nature Methods 11, 1144–1146 (2014).

77. Uritskiy, G. V., DiRuggiero, J. & Taylor, J. MetaWRAP—a flexible pipeline for genome-resolved metagenomic data analysis. Microbiome 6, 158 (2018).

78. Olm, M. R., Brown, C. T., Brooks, B. & Banfield, J. F. dRep: a tool for fast and accurate genomic comparisons that enables improved genome recovery from metagenomes through de-replication. The ISME Journal 11, 2864–2868 (2017).

79. Chklovski, A., Parks, D. H., Woodcroft, B. J. & Tyson, G. W. CheckM2: a rapid, scalable and accurate tool for assessing microbial genome quality using machine learning. Nat Methods 20, 1203–1212 (2023).

80. Chaumeil, P.-A., Mussig, A. J., Hugenholtz, P. & Parks, D. H. GTDB-Tk v2: memory friendly classification with the genome taxonomy database. Bioinformatics 38, 5315–5316 (2022).

81. Almeida, A. et al. A new genomic blueprint of the human gut microbiota. Nature 568, 499–504 (2019).

82. Watson, M. New insights from 33,813 publicly available metagenome-assembled-genomes (MAGs) assembled from the rumen microbiome. 2021.04.02.438222 Preprint at 10.1101/2021.04.02.438222 (2021).

83. Woodcroft, B. J. CoverM. (2023).

84. Woodcroft, B. J., et al. SingleM and Sandpiper: Robust Microbial Taxonomic Profiles from Metagenomic Data. http://biorxiv.org/lookup/doi/10.1101/2024.01.30.578060<x> (</X>2024) doi:10.1101/2024.01.30.578060.

85. Oksanen, J. et al. Vegan: community ecology package. R Package Version 2. 4-6 (2018).

86. Gruber-Vodicka, H. R., Seah, B. K. B. & Pruesse, E. PhyloFlash: Rapid Small-Subunit rRNA Profiling and Targeted Assembly from Metagenomes. mSystems 5, e00920–20 (2020).

87. Quast, C. et al. The SILVA ribosomal RNA gene database project: improved data processing and web-based tools. Nucleic Acids Research 41, D590–D596 (2013).

88. Buchfink, B., Xie, C. & Huson, D. H. Fast and sensitive protein alignment using DIAMOND. Nature Methods 12, 59–60 (2015).

89. Lappan, R. et al. Molecular hydrogen in seawater supports growth of diverse marine bacteria. Nature Microbiology 8, 581–595 (2023).

90. Leung, P. M. & Greening, C. Greening lab metabolic marker gene databases. Monash University (2021).

91. Ortiz, M. et al. Multiple energy sources and metabolic strategies sustain microbial diversity in Antarctic desert soils. Proceedings of the National Academy of Sciences 118, e2025322118 (2021).

92. Deng, Z.-L., Münch, P. C., Mreches, R. & McHardy, A. C. Rapid and accurate identification of ribosomal RNA sequences via deep learning. Nucleic Acids Research 50, e60 (2022).

93. Shen, W., Le, S., Li, Y. & Hu, F. SeqKit: A Cross-Platform and Ultrafast Toolkit for FASTA/Q File Manipulation. PLOS One 11, e0163962 (2016).

94. Robinson, D. broom: An R Package for Converting Statistical Analysis Objects Into Tidy Data Frames. Preprint at 10.48550/arXiv.1412.3565 (2014).

95. Wickham, H. et al. Welcome to the Tidyverse. Journal of Open Source Software 4, 1686 (2019).

96. Wickham, H. Ggplot2: Elegant Graphics for Data Analysis. (Springer, 2016).

97. Poehlein, A. et al. An Ancient Pathway Combining Carbon Dioxide Fixation with the Generation and Utilization of a Sodium Ion Gradient for ATP Synthesis. PLOS One 7, e33439 (2012).

98. Hess, V., Poehlein, A., Weghoff, M. C., Daniel, R. & Müller, V. A Genome-Guided Analysis of Energy Conservation in the Thermophilic, Cytochrome-free Acetogenic Bacterium *Thermoanaerobacter kivui*. BMC Genomics 15, 1139 (2014).

99. Kumar, A. et al. Molecular architecture and electron transfer pathway of the Stn family transhydrogenase. Nat Commun 14, 5484 (2023).

100. Kanehisa, M. & Goto, S. KEGG: Kyoto Encyclopedia of Genes and Genomes. Nucleic acids research 28, 27–30 (2000).

101. Harris, C. R. et al. Array programming with NumPy. Nature 585, 357–362 (2020).

102. Virtanen, P. et al. Fundamental algorithms for scientific computing in python. Nature Methods 17, 261–272 (2020).

103. Seabold, S. & Perktold, J. Statsmodels: econometric and statistical modeling with python. in SciPy vol. 7 (2010).

104. Hunter, J. D. Matplotlib: A 2D graphics environment. Computing in science & engineering 9, 90–95 (2007).

105. Waskom, M. L. seaborn: statistical data visualization. Journal of Open Source Software 6, 3021 (2021).

106. Patro, R., Duggal, G., Love, M. I., Irizarry, R. A. & Kingsford, C. Salmon Provides Fast and Bias-Aware Quantification of Transcript Expression. Nature Methods 14, 417–419 (2017).

107. owkin/PyDESeq2. Owkin (2024).

108. Capella-Gutiérrez, S., Silla-Martínez, J. M. & Gabaldón, T. trimAl: a tool for automated alignment trimming in large-scale phylogenetic analyses. Bioinformatics 25, 1972–1973 (2009).

109. Minh, B. Q. et al. IQ-TREE 2: New Models and Efficient Methods for Phylogenetic Inference in the Genomic Era. Molecular Biology and Evolution 37, 1530–1534 (2020).

110. Letunic, I. & Bork, P. Interactive Tree of Life (iTOL) v6: recent updates to the phylogenetic tree display and annotation tool. Nucleic Acids Research 52, W78–W82 (2024).

111. Asnicar, F. et al. Precise phylogenetic analysis of microbial isolates and genomes from metagenomes using PhyloPhlAn 3.0. Nature Communications 11, 2500 (2020).

112. Segata, N., Börnigen, D., Morgan, X. C. & Huttenhower, C. PhyloPhlAn is a new method for improved phylogenetic and taxonomic placement of microbes. Nature Communications 4, 2304 (2013).

113. Katoh, K. & Standley, D. M. MAFFT Multiple Sequence Alignment Software Version 7: Improvements in Performance and Usability. Molecular Biology and Evolution 30, 772– 780 (2013).

114. Katoh, K., Misawa, K., Kuma, K. I. & Miyata, T. MAFFT: A novel method for rapid multiple sequence alignment based on fast Fourier transform. Nucleic Acids Research 30, 3059–3066 (2002).

115. Jumper, J. et al. Highly accurate protein structure prediction with AlphaFold. Nature 596, 583–589 (2021).

116. Evans, R. et al. Protein complex prediction with AlphaFold-Multimer. Preprint at 10.1101/2021.10.04.463034 (2021).

117. Madhavi Sastry, G., Adzhigirey, M., Day, T., Annabhimoju, R. & Sherman, W. Protein and ligand preparation: parameters, protocols, and influence on virtual screening enrichments. J Comput Aided Mol Des 27, 221–234 (2013).

118. Jarrell, K. F. & Sprott, G. D. The transmembrane electrical potential and intracellular pH in methanogenic bacteria. Can. J. Microbiol. 27, 720–728 (1981).

119. Halgren, T. A. et al. Glide:D A New Approach for Rapid, Accurate Docking and Scoring. 2. Enrichment Factors in Database Screening. Journal of Medicinal Chemistry 47, 1750–1759 (2004).

120. Fischer, A., Smieško, M., Sellner, M. & Lill, M. A. Decision Making in Structure-Based Drug Discovery: Visual Inspection of Docking Results. J. Med. Chem. 64, 2489–2500 (2021).

