## Supplementary Information for "Methanogenesis inhibition remodels microbial fermentation and stimulates acetogenesis in ruminants"

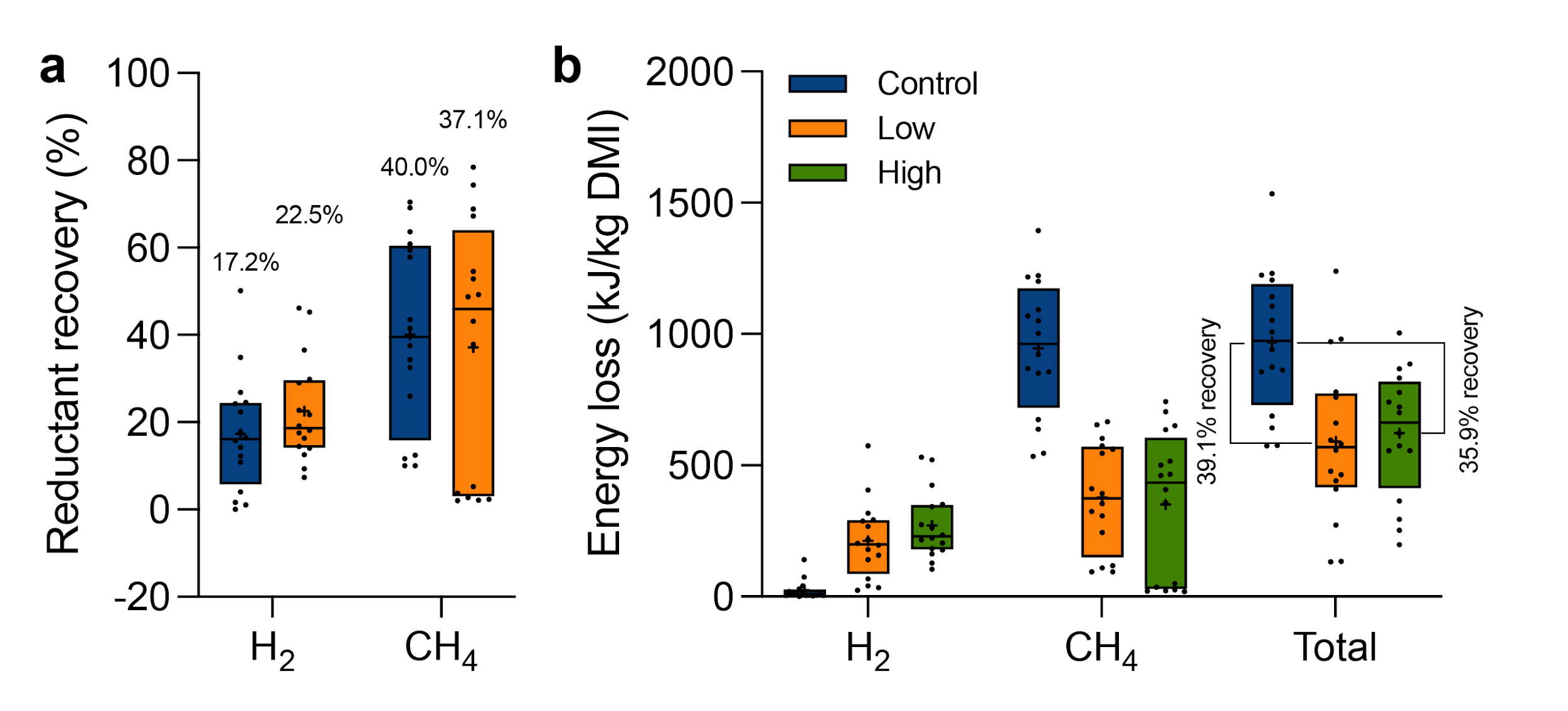

**Figure S1:** Reductant recovery (**a**) and energy loss (**b**) inferred from H_2_ and CH_4_ emissions. (**a**): Reductant recovery indicates the fraction of reducing equivalents lost due to enhanced H_2_ emission and reduced CH_4_ emissions in 3-NOP treated calves relative to the amount of reducing equivalents lost due to CH_4_ emission in the control group. Arithmetic mean values were annotated for each box. (**b**): Energy loss due to the emission of H_2_ and CH_4_.

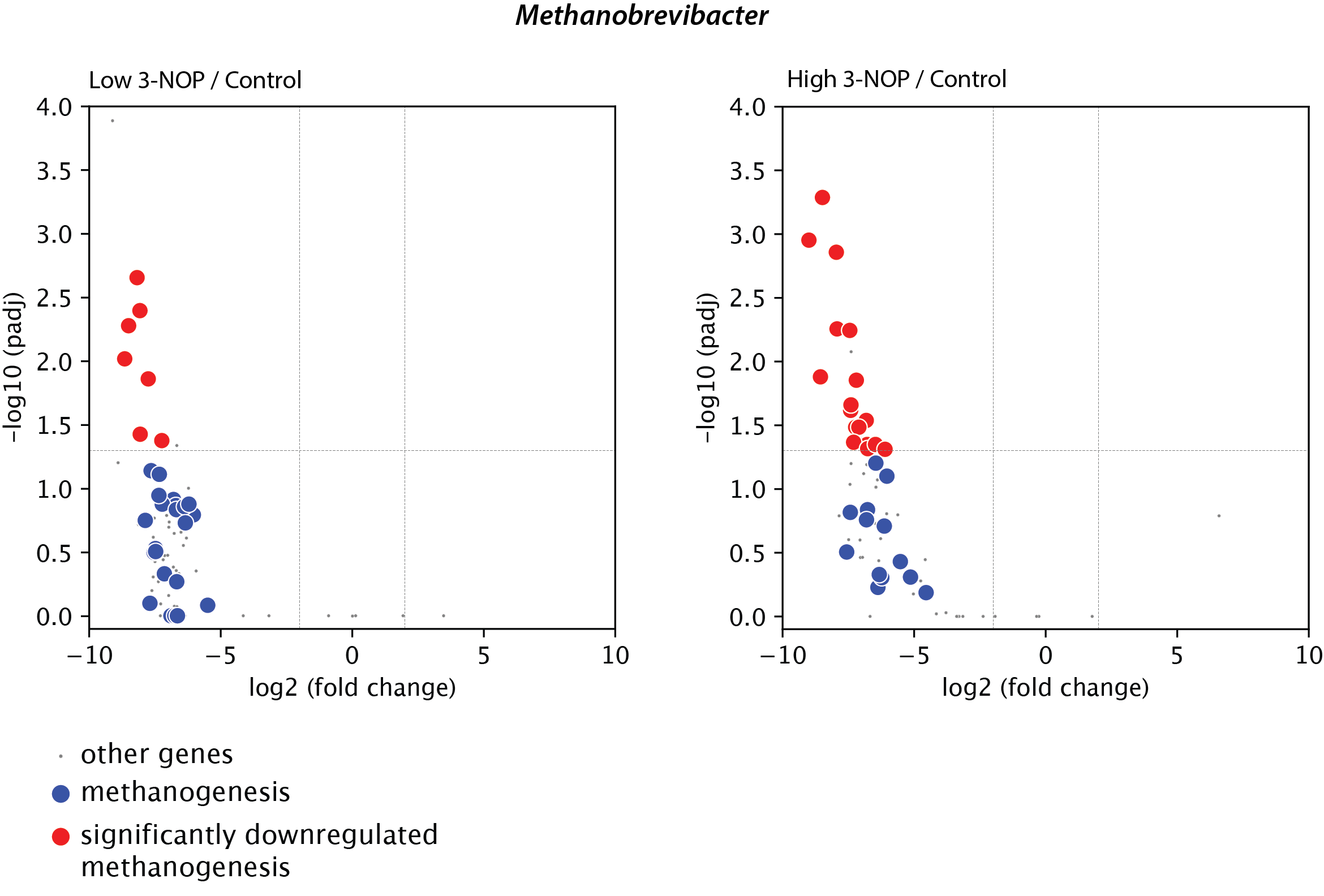

**Figure S2**: Volcano plots of the ratio of transcripts for *Methanobrevibacter* under low 3-NOP versus control (left) and high 3-NOP versus control (right). Each gene is represented by a grey dot; genes from the methanogenesis pathway are indicated by blue dots if their differential expression is insignificant (adjusted p ≥ 0.05), or by red dots when significant. The horizontal dotted line indicates an adjusted p value of 0.05, and the vertical dotted line indicates twofold changes.

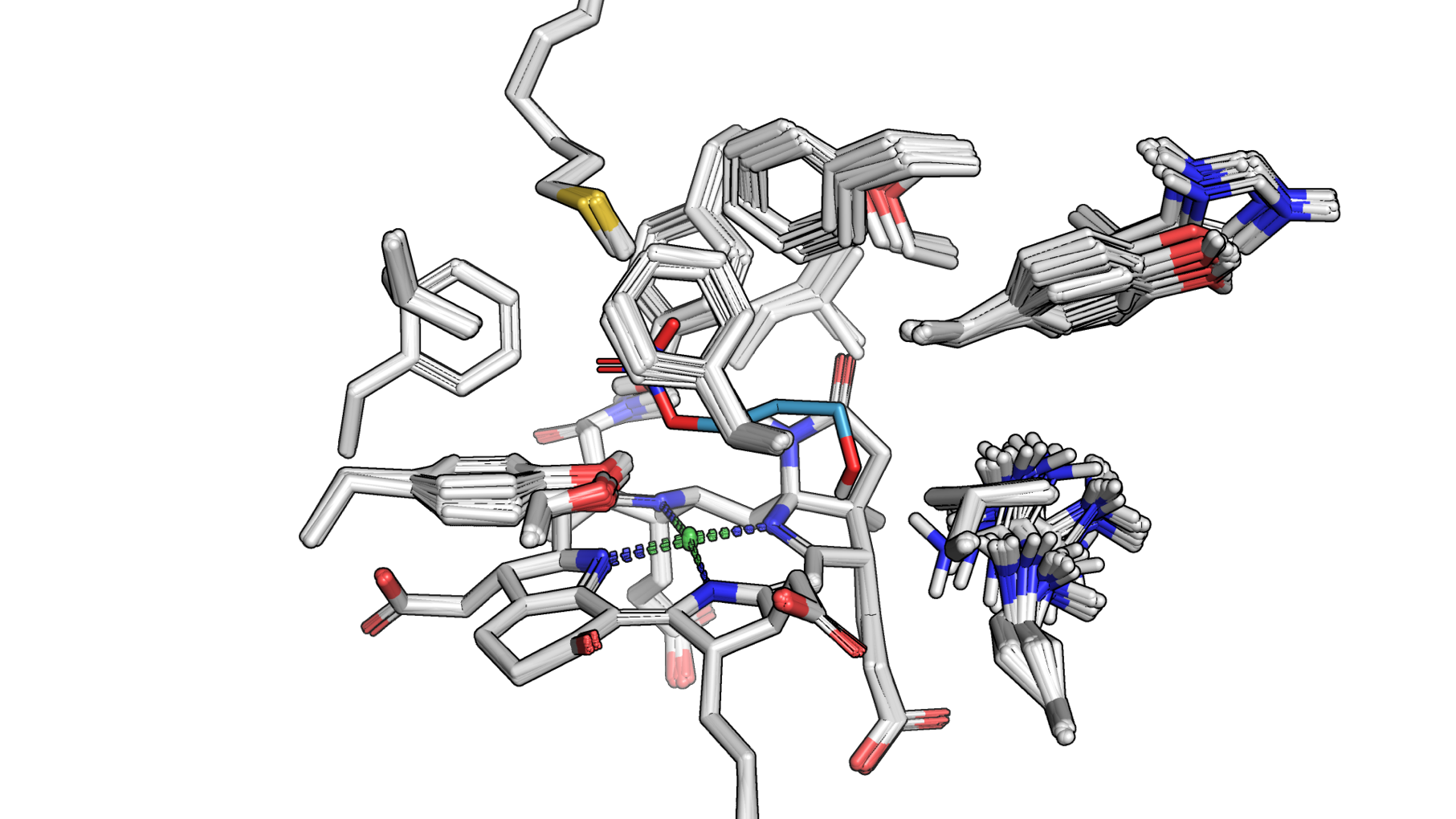

**Figure S3**: A snapshot displaying 3-NOP binding to the active site across 64 superimposed structures of the MCR complex, modelled using AlphaFold2. These protein sequences were predicted from the 49 MAGs analysed in this study. The active site residues, 3-NOP and the F_430_ cofactor are shown in stick representation with carbon, nitrogen, oxygen, and sulfur atoms coloured in white (cyan for 3-NOP), blue, red, and yellow, respectively. The nickel ion of the F_430_ cofactor is shown as a green sphere.

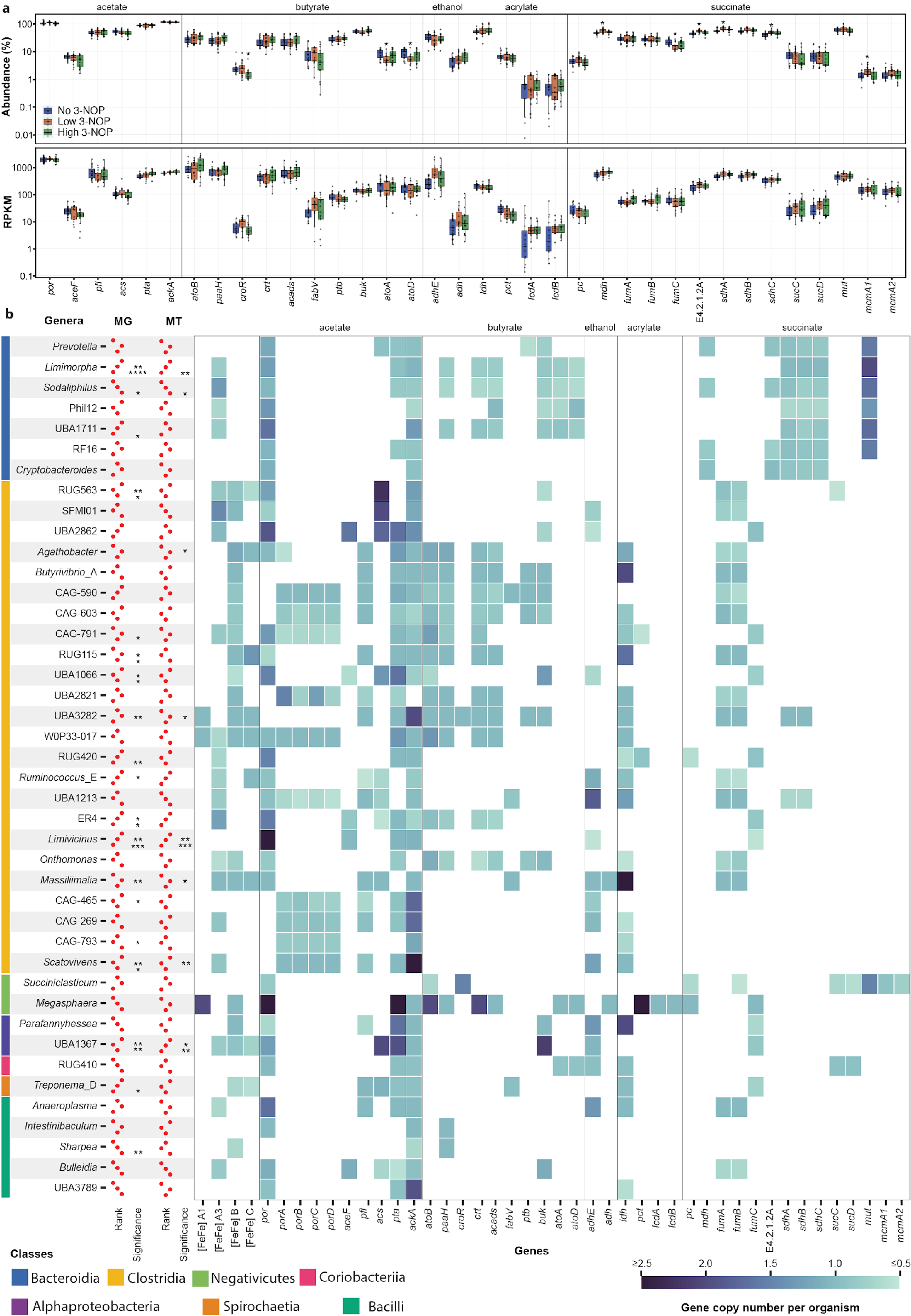

**Figure S4**: (**a**) Abundance and expression of key genes involved in the fermentative formation of acetate, ethanol, butyrate and propionate. Statistical significance of differences between groups (low versus control and high versus control) for each parameter was assessed using the Kruskal–Wallis method with a false discovery rate correction (*p < 0.05). (**b**) Rank abundance of fermentative taxa indicated by dots to show the mean abundance change from control, low and high 3-NOP dosages from top to bottom, and low to high abundance from left to right, in metagenomes and metatranscriptomes. Statistical significance of differences between groups (low versus control and high versus control) for MAG was determined using the Mann–Whitney’s *U* test (* *p* < 0.05, ** *p* < 0.01, *** *p* < 0.001, and **** *p* < 0.0001). A consolidated list of 43 fermentative genera was derived from the 30 most abundant genera under no, low and high 3-NOP conditions. The heatmap displays the average copy numbers of group A1, A3, and B [FeFe]-hydrogenases, alongside genes mediating fermentation pathways for acetate, butyrate, propionate and ethanol production encoded by these organisms.

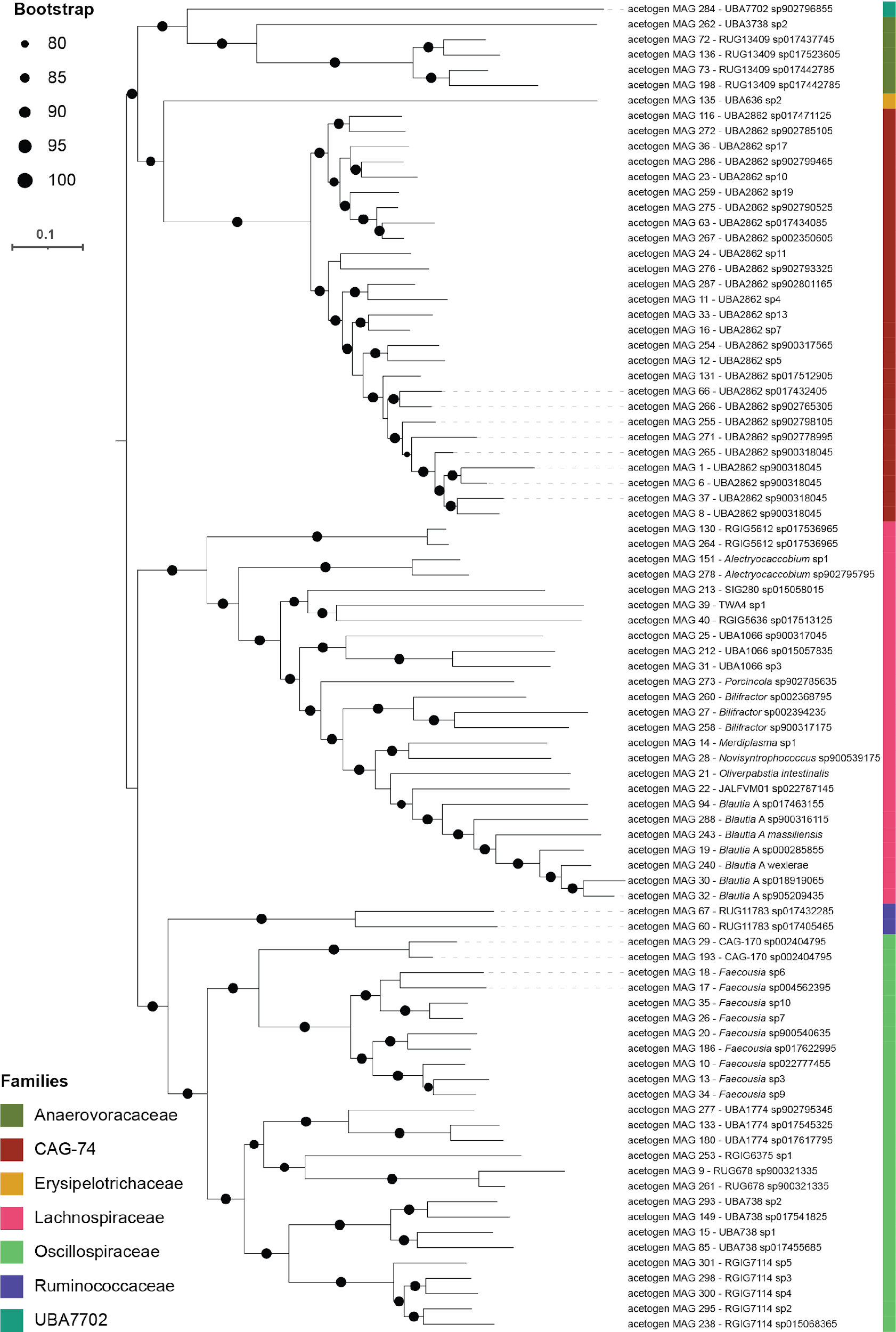

**Figure S5**: Phylogenomic tree illustrating the evolutionary relationships of the acetogen MAGs identified in this study. The tree was inferred using the maximum likelihood method (LG+F+G4) with 1,000 bootstrap iterations and rooted at midpoint.

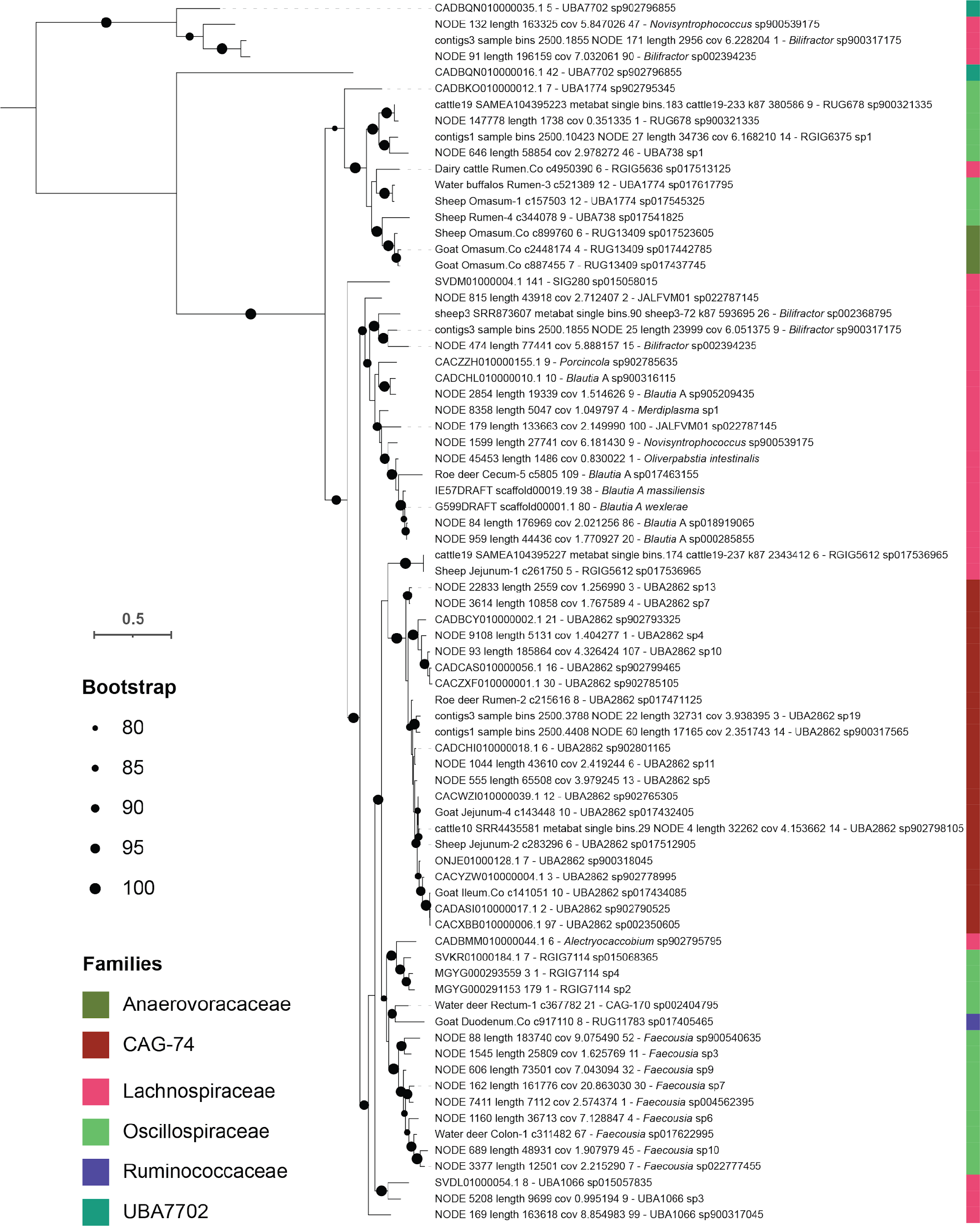

**Figure S6**: Phylogenetic tree consisting of 76 *cooS* (anaerobic carbon monoxide dehydrogenase catalytic subunit) encoded by the acetogens MAGs identified in this study. The tree was inferred using maximum likelihood (LG+I+G4) with 1,000 bootstrap iterations after selecting the best evolutionary model. The tree was rooted at the midpoint. Colour shades on the tree indicate the families from which the protein sequences were sourced.

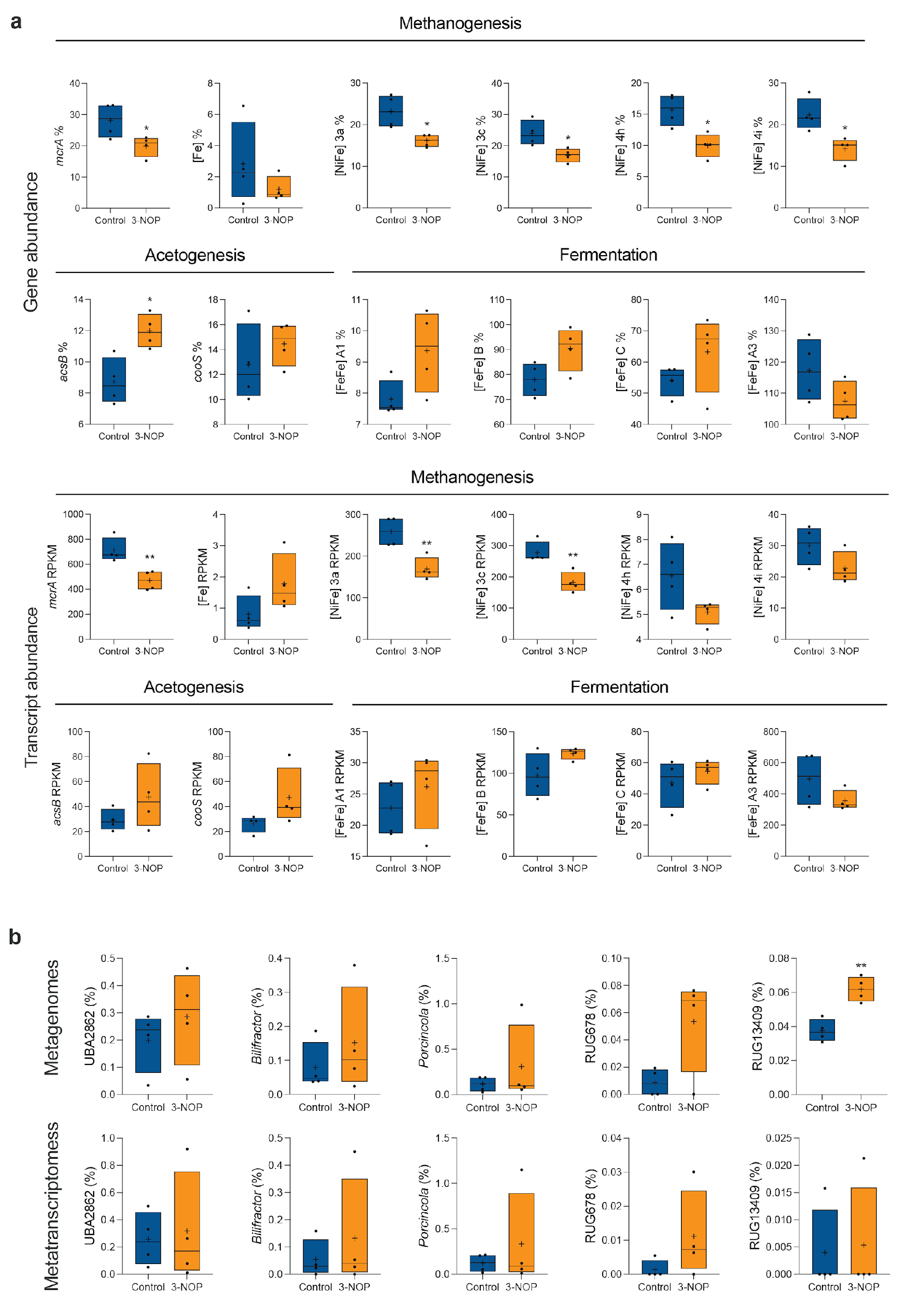

**Figure S7**: Reanalysis of Holstein dairy cattle study^1,2^. (**a**) the abundance (top) and expression (bottom) of metabolic marker genes and hydrogenases involved for methanogenesis, acetogenesis, and fermentation. (**b**) The abundance (top) and activity (bottom) of top-five acetogen genera, analysed through mapping against metagenomes and metatranscriptomes. Statistical significance of differences between groups (low versus control and high versus control) for was assessed using student’s t test (* *p* < 0.05, ** *p* < 0.01).

**References**:

1.	Pitta, D. W. *et al.* The effect of 3-nitrooxypropanol, a potent methane inhibitor, on ruminal microbial gene expression profiles in dairy cows. *Microbiome* **10**, 146 (2022).

2.	Melgar, A. *et al.* Effects of 3-nitrooxypropanol on rumen fermentation, lactational performance, and resumption of ovarian cyclicity in dairy cows. *Journal of Dairy Science* **103**, 410–432 (2020).
